# Central tolerance is impaired in the middle-aged thymic environment

**DOI:** 10.1101/2022.01.17.476690

**Authors:** Jessica N. Lancaster, Damaris L. Keatinge-Clay, Jayashree Srinivasan, Yu Li, Hilary J. Selden, Seohee Nam, Ellen R. Richie, Lauren I. R. Ehrlich

## Abstract

One of the earliest hallmarks of immune aging is thymus involution, which not only reduces the number of newly generated and exported T cells, but also alters the composition and organization of the thymic microenvironment. Thymic T-cell export continues into adulthood, yet the impact of thymic involution on the quality of newly generated T-cell clones is not well established. Notably, the number and proportion of medullary thymic epithelial cells (mTECs) and expression of tissue restricted antigens (TRAs) decline with age, suggesting the involuting thymus may not promote efficient central tolerance. Here, we demonstrate that the middle-aged thymic environment does not support rapid motility of medullary thymocytes, potentially diminishing their ability to scan antigen presenting cells that display the diverse self-antigens that induce central tolerance. Consistent with this possibility, thymic slice assays reveal that the middle-aged thymic environment does not support efficient negative selection or regulatory T cell (Treg) induction of thymocytes responsive to either TRAs or ubiquitous self-antigens. This decline in central tolerance is not universal, but instead impacts lower-avidity self-antigens that are either presented at low levels or bind to TCRs with moderate affinities. Additionally, the decline in thymic tolerance by middle-age is accompanied by both a reduction in mTECs and hematopoietic antigen presenting cell subsets that cooperate to drive central tolerance. Thus, age-associated changes in the thymic environment result in impaired central tolerance against moderate avidity self-antigens, potentially resulting in export of increasingly autoreactive naive T cells, with a deficit of Treg counterparts by middle age.

## Introduction

Thymus involution begins in childhood, resulting in a progressive reduction in the generation and export of naive T cells with age (Chinn et al. 2012). Diminished thymic output contributes to the age-associated decline in T-cell immunity, one of the major drivers of immune dysfunction in aged mice and humans (Elyahu & Monsonego 2021; Nikolich-Žugich 2014). However, the thymus continues to produce and export new T cells, albeit at reduced numbers, into advanced age (Hale et al. 2006; Lynch et al. 2009; Flores et al. 1999). In fact, measurement of human T-cell receptor excision circles (TRECs) indicates that thymic output is detectable until ∼80 years of age (Mitchell et al. 2010; Nasi et al. 2006). Although mature naïve T cells in humans are largely maintained by homeostatic proliferation (Mold et al. 2019), thymic output is required to sustain a normal number of naive T cells in both mice and humans (Appay & Sauce 2014; Bourgeois et al. 2008). Moreover, the thymus remains the sole source of new conventional T-cell and Treg clones throughout life, yet little is known about the impact of thymus aging on qualitative changes in T-cell maturation and selection.

As the thymus involutes and fewer recent thymic emigrants are exported, the peripheral T compartment is progressively comprised of cells with a memory phenotype (Nikolich-Žugich 2008; Srinivasan et al. 2021; Goronzy & Weyand 2019). This reduction in naïve T cells in the elderly contributes to increased susceptibility to infectious disease and decreased responsiveness to vaccines (Nikolich-Žugich 2014). Interestingly, while the number of new T cells exported from the thymus declines substantially by middle-age (Mold et al. 2019; den Braber et al. 2012; Ito et al. 2017), the incidence of new-onset autoimmunity peaks at middle-age for many autoimmune disorders (Watad et al. 2017). Both naive CD4^+^ and CD8^+^ T cells (Quinn et al. 2016; Rudd et al. 2011; Deshpande et al. 2015) become more self-reactive with age in mice, suggesting that changes in thymic selection could contribute to increased T-cell auto-reactivity with age.

The thymic medulla is a specialized microenvironment for inducing T-cell central tolerance to diverse autoantigens. Following T-lineage commitment, differentiation, and positive selection in the thymic cortex, developing T cells express chemokine receptors that promote their entry into the medulla (Cowan et al. 2014; Kadakia et al. 2019; Kurobe et al. 2006; Lancaster et al. 2018; Ehrlich et al. 2009; Hu et al. 2015), where they encounter numerous self-antigens presented by medullary thymic epithelial cells (mTECs) and hematopoietic antigen presenting cells (HAPCs), including conventional dendritic cells (cDCs), B cells, and plasmacytoid dendritic cells (pDCs). If a thymocyte expresses a T-cell receptor (TCR) with sufficiently high affinity for these self-antigens, it undergoes negative selection or diversion to the Treg lineage, enforcing central tolerance (Klein et al. 2014). Mature mTECs play an essential role in tolerance induction, as they collectively express about 90% of the proteome, including *Aire-*dependent tissue-restricted antigens (TRAs), which are otherwise expressed in only a few peripheral tissues (Bautista et al. 2021; Bornstein et al. 2018; Brennecke et al. 2015; Meredith et al. 2015; Sansom et al. 2014). It is critical that thymocytes are tolerized to the full repertoire of mTEC-derived self-antigens to avoid autoimmunity (Aaltonen et al. 1997; Anderson 2002; DeVoss et al. 2006; Nagamine et al. 1997), but a given TRA is expressed by only ∼1-3% of mTECs (Derbinski et al. 2005; Derbinski et al. 2008), creating a sparse mosaic of self-antigen display in the medulla. Thymic cDCs also play a critical role in thymic tolerance by presenting self-antigens acquired from circulation, peripheral tissues, and mTECs (Bonasio et al. 2006; Atibalentja et al. 2011; Perry et al. 2014, 2018; Leventhal et al. 2016; Watanabe et al. 2020; Vollmann et al. 2021). Thus, it is critical that post-positive selection thymocytes enter the medulla efficiently and rapidly scan mTECs and HAPCs to encounter the complete arrays of self-antigens needed to induce broad central tolerance.

Age-associated changes in thymic APCs and self-antigen expression could impair the ability of the thymus to support central tolerance. Aging is associated with substantial changes in thymic stromal organization and composition (Baran-Gale et al. 2020; Chinn et al. 2012; Lynch et al. 2009; Srinivasan et al. 2021; Venables et al. 2019). In the aged thymus, the cortex thins (Venables et al. 2019), TEC proliferation and cellularity is greatly reduced (Gray et al. 2006), and the frequency and number of mTECs decline (Chinn et al. 2012; Lepletier et al. 2019; Baran-Gale et al. 2020). Notably, expression of TRAs in the medulla diminishes with age (Griffith et al. 2012), and thymic B cells and DCs change in composition and molecular properties (Cepeda et al. 2018; Flores et al. 2001; Nuñez et al. 2016; Ki et al. 2014; van Dommelen et al. 2010; Varas et al. 2003).

The efficiency with which auto-reactive thymocytes are negatively selected is dependent on the TCR-binding avidity of the selecting self-antigen. High avidity self-antigens induce efficient negative selection (Klein et al. 2019), while thymocytes expressing TCRs that bind self-antigen with moderate avidity can escape negative selection, resulting in autoimmune pathology (Koehli et al. 2014; Zehn & Bevan 2006). Beyond TCR binding affinity, the pattern of self-antigen expression in the thymus also modulates autoreactive thymocyte fates. Ubiquitously-expressed self-antigens tend to induce more robust negative selection, while rare, *Aire*-dependent TRAs induce both negative selection and Treg induction (Malhotra et al. 2016; Hassler et al. 2019). Given that aging results in diminished expression of TRAs and changes in the composition and organization of mTECs and HAPCs, autoreactive thymocytes in the aged microenvironment may be screened less efficiently against ubiquitous and/or rare, tissue-specific self-antigens. However, little is known about the impact of aging on thymocyte selection. In a genetically-induced mouse model of accelerated thymic involution, negative selection was impaired, but Treg induction was enhanced (Coder et al. 2015; Oh et al. 2017). In contrast, following natural aging, generation of new thymic Tregs diminished (Thiault et al. 2015), which was attributed to an increase in peripheral Treg re-entering the thymus and outcompeting resident Treg progenitors for IL-2 (Hemmers et al. 2019; Weist et al. 2015). These findings raise questions about whether the aged thymus supports efficient negative selection and Treg induction against different types of self-antigens.

In this study, we use live thymic slices in combination with 2-photon microscopy (2PM) to test the ability of the naturally aged thymic environment to support thymocyte medullary entry and rapid motility, as well as negative selection and Treg induction in response to ubiquitous self-antigens or model TRAs. We find that thymocytes, regardless of age, do not migrate as rapidly in a middle-aged 12-month (MO) relative to a 1MO thymic environment. Furthermore, the middle-aged thymus does not support efficient negative selection or Treg induction of thymocytes responsive to self-antigens of moderate avidities. However, central tolerance, including Treg induction, remains intact for thymocytes responsive to ubiquitous high-affinity self-antigens in the middle-aged thymus. Thus, the middle-aged thymus does not support efficient central tolerance to moderate-avidity self-antigens, possibly resulting in export of poorly tolerized T cells to the periphery by middle-age.

## Results

### The middle-aged thymus environment does not support rapid motility of medullary thymocytes

To test whether the middle-aged thymic environment supports rapid motility and efficient accumulation of post-positive selection thymocytes in the medulla, we used 2PM to image young 1MO and middle-aged 12MO polyclonal mouse CD4^+^ single-positive (CD4SP) thymocytes migrating in either 1MO or 12MO live thymic slices (**Figure 1a**). We first determined that expression of CCR4 and CCR7, which promote medullary entry (Ehrlich et al. 2009; Hu et al. 2015; Ueno et al. 2004), are comparable between 1MO and 12MO thymocytes (**Supplementary Figure 1a**). Next, CD4SPs from 1MO and 12MO thymuses were labeled with red or blue fluorescent dyes and allowed to migrate in 1MO versus 12MO thymic slices generated from pCX-EGFP mice, in which cortical and medullary regions can be distinguished by cellular morphology and EGFP intensity (Lancaster & Ehrlich 2017). Thymocyte migration was imaged by time-lapse 2PM, and cells were tracked to determine their density in the medulla and cortex, as well their velocity and path straightness (**Figure 1b and Supplementary Movies 1-2)**.

**Figure 1.**
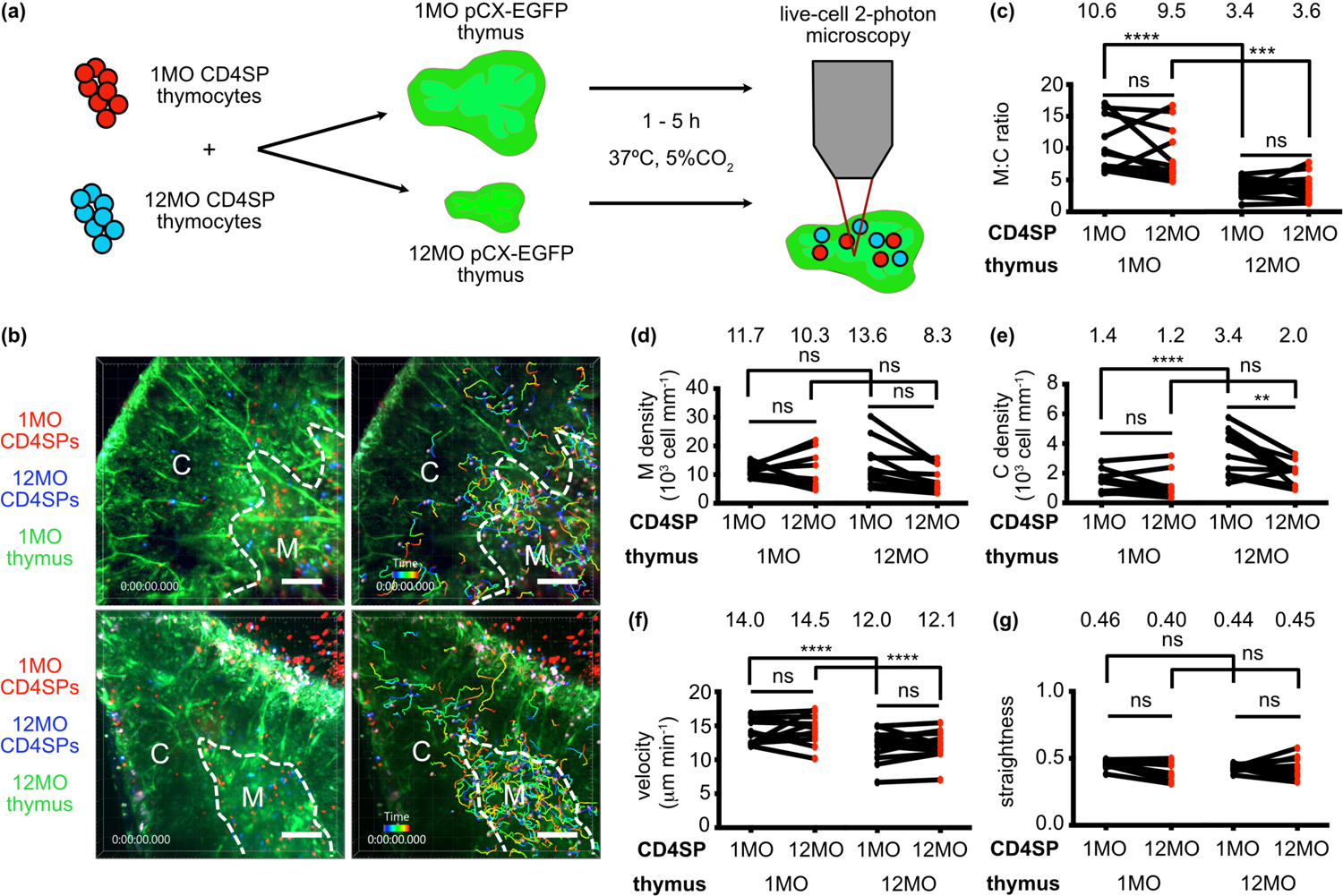
The middle-aged thymus environment does not support rapid motility of medullary thymocytes. **(a)** Schematic of 2PM approach for imaging migration of CMPTX (red)- or Indo1AM (blue)-labeled 1MO vs 12MO polyclonal CD4SP thymocytes in 1MO or 12MO live pCX-EGFP (green) thymic slices. Time-lapse imaging through a 40-μm depth was carried out for 15 minutes to visualize CD4SP localization and migratory properties. **(a)** Representative maximum intensity projections of 2PM imaging volumes at 20X magnification. The cortex (C) and medulla (M) are delineated by dashed white lines. 1MO (red) and 12MO (blue) CD4SP cells can be seen in the left images, while the right images show the same images with cell tracks color-encoded for elapsed imaging time. Scale bars, 100 μm. **(c-e)** Quantification of the density of 1MO and 12MO CD4SP cells in the **(b)** medullary versus cortical imaging volumes (M:C ratio), **(d)** the medulla and **(e)** the cortex of 1MO vs 12MO thymic slices. **(f)** Mean cell velocity and **(g)** track straightness of 1MO and 12MO thymocytes migrating within 1MO versus 12MO thymic slices. Data are compiled from 4 experiments, with each point indicating the mean thymocyte value within a given thymic slice (*n*_1MO_ = 11, *n*_12MO_ = 12). Total cells tracked: *n*_1MO_ = 903 and *n*_12MO_ = 607 in 1MO slices, *n*_1MO_ = 1027 and *n*_12MO_ = 586 in 12MO slices. Analyzed by *t*-tests, *p*-values: ** < 0.01, *** < 0.001, **** < 0.0001, ns: not significant. See also Supplementary Figure 1 and Movies 1 and 2.

The ratio of CD4SPs within the medulla relative to the cortex declined in the 12MO versus 1 MO thymic environment, irrespective of thymocyte age (**Figure 1c**). However, the density of CD4SP cells in the medulla of 12MO thymuses was not significantly reduced (**Figure 1d**); instead, the density of CD4SP cells in the 12MO cortex increased, likely reflecting age-associated cortical thinning (Venables et al. 2019; Chinn et al. 2012) (**Figure 1e**). Consistent with robust accumulation of CD4SP in the12MO medulla, expression of CCL21, the CCR7 ligand required for accumulation of CD4SP thymocytes in the medulla (Kozai et al. 2017), was elevated in the 12MO versus 1MO medulla (**Supplementary Figure 1b**). Notably, CD4SPs of both ages migrated significantly more slowly in the 12MO versus 1MO thymus (**Figure 1f**). Neither the age of the thymocytes nor the thymic environment significantly impacted the straightness of thymocyte migration paths (**Figure 1g**). Thus, the middle-aged thymus environment supports robust entry of CD4SP cells into the medulla but does not support their rapid migration, both of which enable efficient scanning of self-antigens presented by medullary APCs.

### Thymic slice deletion assays for quantification of age-associated changes in central tolerance

We next investigated whether central tolerance induction is impaired in the middle-aged thymic environment using live thymic slice deletion assays. Negative selection of young 1MO CD8SP or CD4SP thymocytes responding to self-antigens in 1MO versus 12MO thymic slices was quantified by flow cytometry (**Figure 2a**) (Hu et al. 2015; Lancaster et al. 2019). TCR transgenic thymocytes specific for ovalbumin peptides (OVAp) were mixed at an equal ratio with polyclonal CD45-congenic thymocytes prior to loading onto 1MO or 12MO thymic slices. The polyclonal thymocytes served as non-selecting controls to normalize for differential cell entry into individual slices. To assay for negative selection, the slices 1) lacked ovalbumin (OVA-), serving as negative controls for selection, 2) were incubated with exogenously administered OVAp, modeling ubiquitous self-antigens, or 3) were generated from transgenic mice expressing OVA under control of the *Aire*-dependent rat insulin promoter (RIP), modeling endogenous TRAs (Figure 2A). CD8SP negative selection was assayed with OT-I TCR transgenic thymocytes, which express a TCR specific for OVAp_257-264_ (SIINFEKLp) presented by H-2K^b^ (Hogquist et al. 1994). One advantage of the OT-I system is that altered peptide ligands (APLs) of varying affinities for the OT-I TCR been defined and expressed as model TRAs under control of the RIP (Koehli et al. 2014; Daniels et al. 2006). CD4SP deletion was assayed with OT-II TCR transgenic thymocytes, which recognize OVAp_323-339_ presented by I-A^b^ (Barnden et al. 1998). This slice deletion assay allows us to test the impact of a naturally aged, non-irradiated thymic microenvironment on central tolerance, without potentially confounding differences in thymocyte ages. By varying the concentration and/or TCR-binding affinity of the OVAp APLs added to the slices, the impact of TCR-binding avidity on age-associated changes in negative selection efficiency can also be tested.

**Figure 2.**
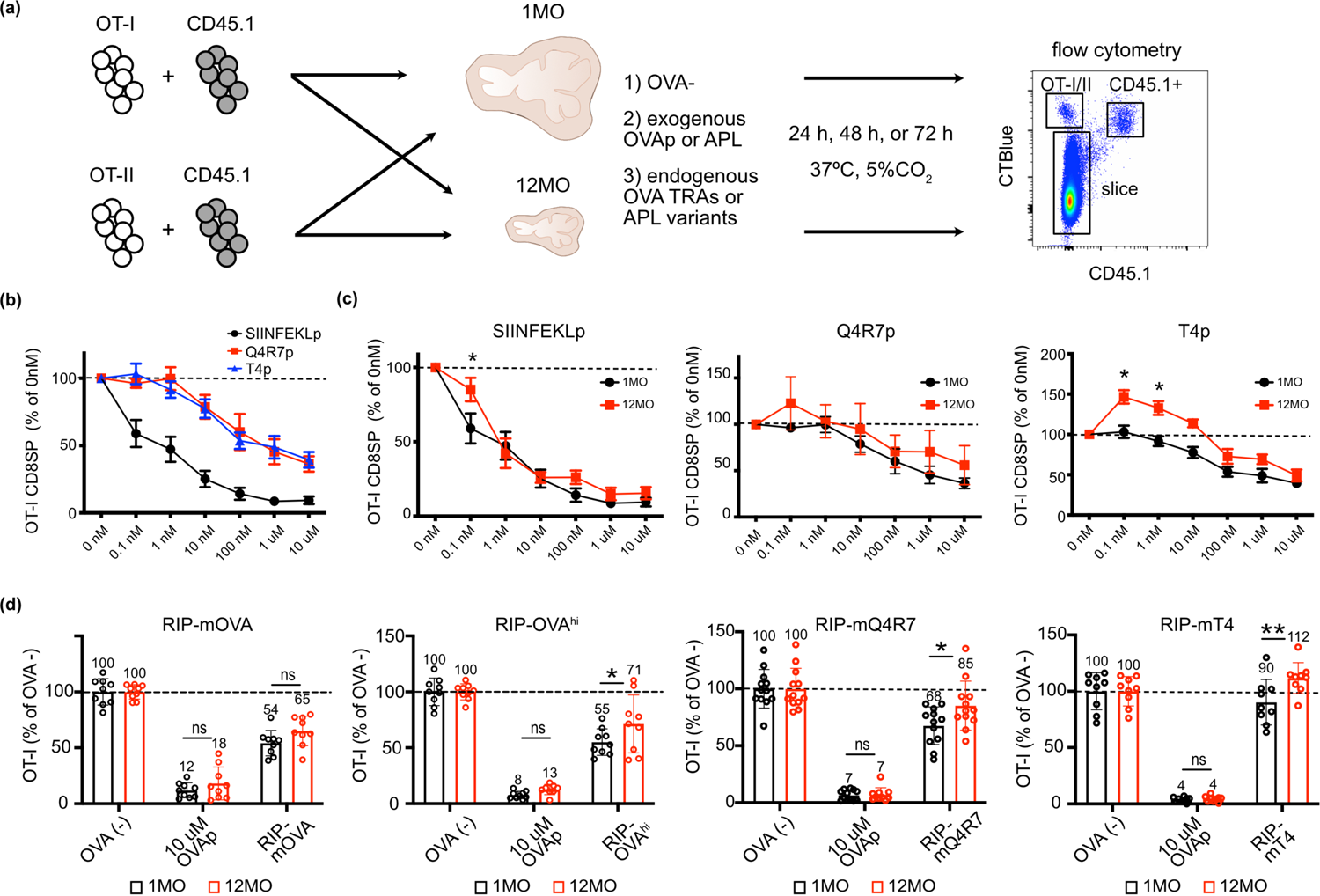
The 12MO thymic environment does not support efficient negative selection of OT-I CD8SP thymocytes responding to moderate avidity self-antigens. **(a)** Experimental approach of heterochronic slice deletion assays to assess negative selection of OT-I thymocytes responding to ubiquitous self-antigens or TRAs in young (1MO) vs middle-aged (12MO) thymic slices. Slices were generated from 1) C57BL/6J wild-type mice that did not express OVA (OVA-), 2) wild-type mice followed by incubation with OVAp or lower affinity APLs, or 3) the TRA strains RIP-mOVA, RIP-OVA^hi^, RIPmQ4R7, or RIPmT4. **(b)** OT-I CD8SP cellularity in 1MO thymic slices incubated overnight with the indicated concentrations of SIINFEKLp, Q4R7p, or T4p, normalized to OT-I CD8SPs in slices incubated without peptide. Data compiled from 6 experiments. **(c)** OT-I CD8SP cellularity on 1MO or 12MO thymic slices incubated with the indicated concentrations of SIINFEKLp, Q4R7p or T4p. Data are normalized to OT-I CD8SPs in slices incubated without peptide. Data compiled from 6-7 independent experiments, with data points representing mean of triplicate thymic tissue slices. Data in **(b)** is a composite of the 1MO data shown in **(c)**. **(d)** Negative selection of OT-I CD8SP thymocytes responding to AIRE-dependent TRAs in 1MO and 12MO thymic slices, evaluated at 48 h. The proportions of OTI CD8SPs remaining in RIP-mOVA, RIP-OVA^hi^, RIP-mQ4R7 and RIP-mT4 thymic slices relative to the numbers in numbers in thymic slices that do not express OVA. Addition of 10 μM of SIINFEKLp (OVAp) served as a positive control for OT-I negative selection. Data show mean ± SEM compiled from 9 independent experiments, with data points representing mean of triplicate thymic tissue slices and values normalized to the mean of triplicate OVA^-^ slices. Data in **(c)** and **(d)** were analyzed by two-way ANOVA with Šídák’s correction for multiple comparisons, *p*-values: * < 0.05, ** < 0.01, ns: not significant.

### The middle-aged thymic environment does not support efficient negative selection of CD8SP thymocytes responding to moderate avidity self-antigens

To test whether the middle-aged thymus environment supports negative selection of CD8SP cells responding to ubiquitous self-antigens, 1MO OT-I thymocytes were introduced onto 1MO vs 12MO thymic slices incubated with varying concentrations of SIINFEKLp or OVA APLs. SIINFEKLp has a relatively high affinity for the OT-I TCR (K_d_ 3.7 ± 0.7nM) compared with Q4R7p (K_d_ 48 ± 9.5 nM) and T4p (K_d_ 55 ± 10.1 nM), (Daniels et al. 2006). All three peptides induced negative selection in a concentration dependent manner after 24hr on both 1MO and 12MO thymic slices, and negative selection efficiency correlated with the affinity for the OT-I TCR (**Figure 2b**). Notably, at higher peptide concentrations, there was negligible difference in the extent of CD8SP negative selection in 1MO versus 12MO thymus environments (**Figure 2c**). However, deletion on 12MO slices was significantly diminished at lower concentrations of SIINFEKLp and T4p, with a similar trend for Q4R7p (**Figure 2c**). In response to the low affinity T4p, the number of CD8SPs in 12MO slices pulsed with 0.1-10 nM T4p exceeded that of the no-peptide control, likely indicating a switch to positive selection in the presence of low concentrations of a weak agonist in the middle-aged thymus environment. Together, these results indicate that the middle-aged thymic microenvironment becomes impaired in its ability to support efficient negative selection of CD8SP thymocytes responding to low avidity ubiquitous self-antigens.

To determine whether the middle-aged thymic environment supports negative selection of OT-I thymocytes responding to endogenous TRAs, thymic slices were generated from RIP-mOVA or RIP-OVA^hi^ mice, expressing membrane-bound or soluble forms of OVA, respectively (Kurts et al. 1996; Kurts et al. 1998). These model TRAs induce OT-I CD8SP negative selection *in vivo* (Gallegos & Bevan 2004; Hubert et al. 2011) and in thymic slices (Lancaster et al. 2019). Thymic slices from OVA^-^ littermates incubated with or without 10 μM OVAp served as negative and positive controls for deletion, respectively. Relative to 1MO thymic slices, the 12MO thymic environment supported comparable deletion of OT-I thymocytes to RIP-mOVA. However, negative selection was significantly impaired in middle-aged thymic slices when OT-I thymocytes responded to the RIP-OVA^hi^ TRA (**Figure 2d**), which is expressed at lower levels than the RIP-mOVA TRA (Lancaster et al. 2019). In addition, the 12MO thymic environment did not support efficient negative selection of OT-I thymocytes responding to lower affinity RIP-mQ4R7 and RIP-mT4 (**Figure 2d**). Together, these data indicate that negative selection is impaired in the middle-aged thymus for CD8SP thymocytes responding to TRAs with lower TCR-binding avidities, due to either low expression levels or reduced TCR-binding affinities of the selecting self-peptides.

### The middle-aged thymic environment does not support efficient negative selection of CD4SP thymocytes responding to lower avidity self-antigens

To determine if the middle-aged thymus supports efficient negative selection of CD4SP thymocytes, 1MO OT-II thymocytes were introduced into 1MO versus 12MO thymic slices with varying concentrations of OVAp_323-339._ Slices of both ages induced concentration-dependent negative selection of OT-II CD4SPs after 24hr (**Figure 3a**). Notably, 12MO thymic slices were significantly impaired in their ability to support negative selection of OT-II thymocytes to 10μM OVAp as well as the TRAs RIP-mOVA and RIP-OVA^hi^ at 48 hr (**Figure 3b**). Thus, negative selection of CD4SP cells responsive to both ubiquitous self-antigens and TRAs is impaired in the 12MO thymic environment.

**Figure 3.**
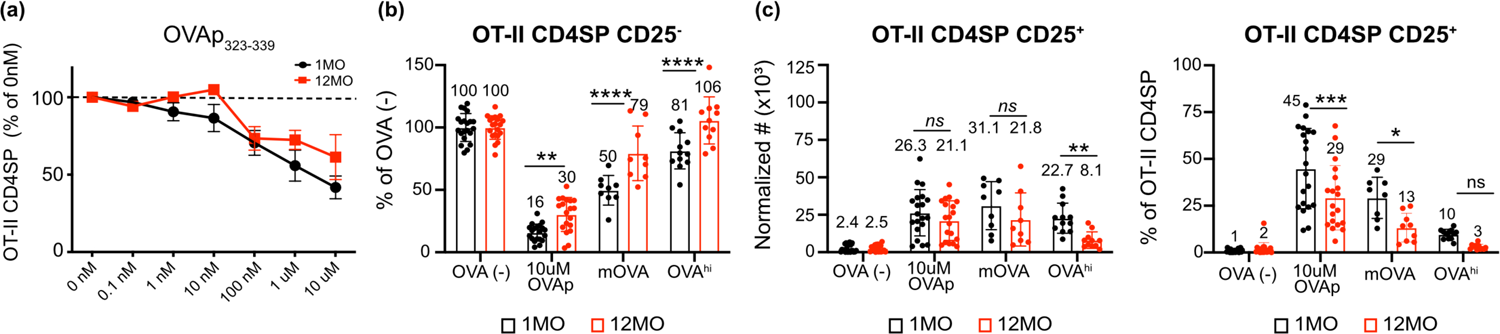
The middle-aged thymic environment does not support efficient negative selection of OT-II CD4SP thymocytes responding to ubiquitous self-antigens or TRAs. (a) OT-II CD4SP cellularity was quantified following incubation overnight in 1MO or 12MO thymic slices with the indicated concentrations of OVAp_323-339_. The frequencies of OT-II CD4SP cells remaining relative to those in slices incubated without OVAp were quantified. Plots show mean ± SEM of compiled data from four independent experiments. Analyzed by two-way ANOVA with Šídák’s correction for multiple comparisons. (b) Negative selection of CD4SP thymocytes and (c) induction of CD25^+^ Treg precursors to the indicated *Aire*-dependent TRAs on 1MO or 12MO thymic slices were quantified. Data show the relative proportions of OT-II CD4SP cells remaining in RIP-mOVA and RIP-OVA^hi^ thymic slices relative to thymic slices that do not express OVA. Addition of 10 μM of OVAp_323-339_ (OVAp) served as a positive control for OT-II negative selection. Data in (b-c) show mean ± SEM compiled from 4 independent experiments, with data points representing mean of triplicate thymic tissue slices. Analyzed by two-way ANOVA with Šídák’s correction for multiple comparisons, *p*-values: * < 0.05, ** < 0.01, *** < 0.001, **** < 0.0001, ns: not significant. See also Supplementary Figure 2.

We also assessed whether diversion of OT-II CD4SP thymocytes towards the Treg lineage was impaired in the 12MO thymic environment by quantifying OT-II CD4SP CD25^+^ Treg precursors (Treg-P) (Hsieh et al. 2004). While the number of Treg-P generated in a middle-aged thymus did not decline significantly in response to exogenous OVAp or the RIP-mOVA TRA after 48hr of selection, the frequency of CD4SP cells upregulating CD25 was significantly diminished in response to both the ubiquitous OVAp self-antigen and the RIP-mOVA TRA (**Figure 3c** and **Supplementary Figure 2a**). These findings indicate that CD4SP cells can be induced, albeit less efficiently, to divert towards a Treg fate when strong self-antigens are presented ubiquitously or as TRAs in the middle-aged thymus. However, in response to the less abundant RIP-OVA^hi^ TRA, the number OT-II CD4SP CD25^+^ Treg-P declined significantly in the 12MO thymic environment, and their frequency declined by over 3-fold (**Figure 3c** and **Supplementary Figure 2a**). Thus, diversion of autoreactive OT-II CD4SPs towards the Treg fate is particularly inefficient in response to lower avidity TRAs in the middle-aged thymic environment.

### Treg induction in response to TRAs is less efficient in the middle-aged thymus

Previous studies showed that by 12MO of age, the mouse thymus generates very few new Treg (Thiault et al. 2015). Because we found that the 12MO thymic environment supports the induction of OT-II CD25^+^ CD4SP Treg-P in response to ubiquitous self-antigens, but not lower abundance TRAs (**Figure 3**), we wondered whether the age-associated decline in *de novo* differentiation of Tregs preferentially impacts thymocytes responsive to a subset of self-antigens. 1 versus 12MO thymic slices pulsed with exogenous OVAp or expressing the RIP-mOVA or RIP-OVA^hi^ TRAs were tested for their ability to support OT-II CD4SP Treg differentiation after 72hr to enable sufficient time for FOXP3 upregulation (Weist et al. 2015). Input OT-II thymocytes contained virtually no detectable Tregs or Treg precursors (**Supplementary Figure 2b**).

First we tested differentiation of CD25^+^FOXP3^+^ Treg and the two precursor populations, CD25^+^FOXP3^-^ (CD25^+^) Treg-P and CD4^+^CD25^-^FOXP3^lo^ (FOXP3^lo^) Treg-P (Owen et al. 2019) in response to ubiquitous self-antigens (**Figure 4a**). In the absence of cognate antigen (OVA^-^ slices), OT-II thymocytes did not differentiate into Treg-P or Tregs. In the presence of 1μM OVAp_323-339_, middle-aged slices supported generation of CD25^+^ Treg-P and Treg as efficiently as young slices, although there was a slight decline in the frequency of Treg in 12MO slices (**Figures 4a, b,d**). Notably, ∼60% of OT-II CD4SP thymocytes upregulated CD25 following addition of OVAp, regardless of the thymic microenvironment age (**Figures 4a-b**), indicating that access to ubiquitous antigens is not impaired in the middle-aged thymus. Exogenous OVAp did not induce many FOXP3^lo^ Treg-P (**Figures 4a, c**). Together, these results demonstrate that the middle-aged thymic environment efficiently supports the *de novo* generation of CD25^+^ Treg-P and Treg from thymocytes responding to ubiquitously presented self-antigens.

**Figure 4.**
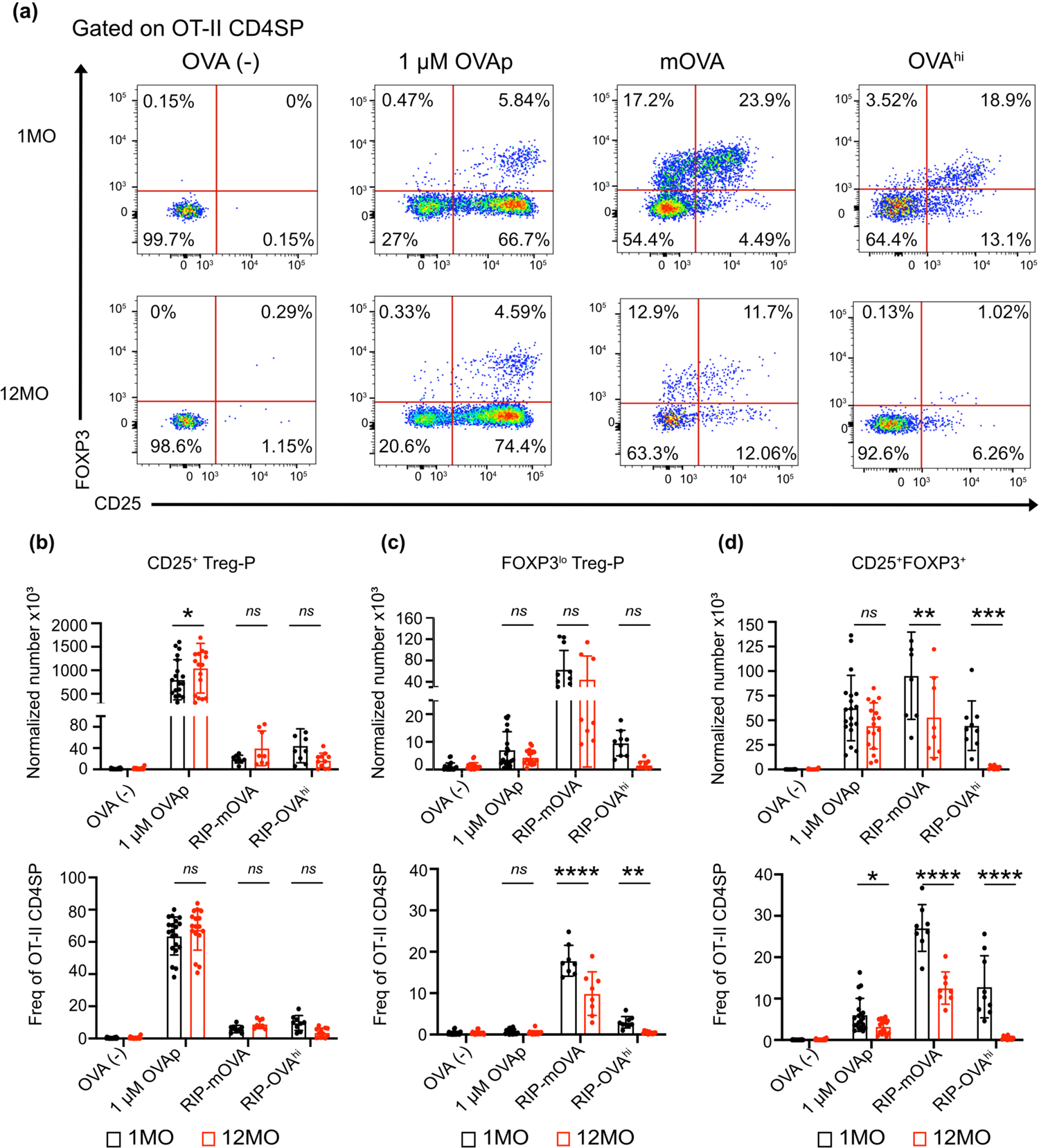
The middle-aged thymic environment does not support efficient induction of Tregs in response to TRAs. **(a)** Representative flow cytometry plots of Treg precursors (CD25^+^ Treg-P and FOXP3^lo^ Treg-P) and Tregs (CD25^+^FOXP3^+^) recovered from OVA(-), RIP-mOVA and RIP-OVA^hi^ 1MO versus 12MO live thymic tissue slices 72 hr post-incubation. Addition of 1 μM of OVAp_323-339_ served as a positive control for Treg induction. Normalized cell numbers (top) and frequencies of OT-II CD4SP cells (bottom) of **(b)** CD25^+^ Treg-P, **(c)** FOXP3^lo^ Treg-P and **(d)** Treg cells. Data in **(b-d)** show mean ± SEM compiled from 3-4 independent experiments, with data points representing mean of triplicate thymic tissue slices. Analyzed by two-way ANOVA with Šídák’s correction for multiple comparisons, *p*-values: * < 0.05, ** < 0.01, *** < 0.001, **** < 0.0001, ns: not significant.

Next, we tested the efficiency of Treg-P differentiation in response to the *Aire*-dependent RIP-mOVA and RIP-OVA^hi^ TRAs in middle-aged thymic slices. 12MO slices supported CD25^+^ Treg-P differentiation in response to TRAs as well as 1MO slices, although many fewer CD25^+^ Treg-P were generated in response to TRAs compared to OVAp on slices of both ages (**Figure 4a-b**). Interestingly, in comparison to the other self-antigens, the RIP-mOVA TRA induced the highest frequency and number of FOXP3^lo^ Treg-P in slices of both ages (**Figures 4c**). Notably, the frequency of FOXP3^lo^ Treg-P declined significantly in the middle-aged thymic environment for both RIP-mOVA and RIP-OVA^hi^ TRAs, with a trend towards decreasing FOXP3^lo^ Treg-P numbers (**Figures 4a,c**). Differentiation of CD25+ FOXP3+ Tregs in response to TRAs was the most significantly impaired in the naturally aged thymic environment. The RIP-mOVA TRA induced efficient Treg generation in 1MO slices, in which ∼25% of remaining OT-II CD4SP thymocytes at 72 h were Tregs. Both the number and frequency of Tregs were markedly lower in middle-aged RIP-mOVA slices, with an almost three-fold decline in frequency in the 12MO versus 1MO thymus (**Figure 4d**). An even more dramatic decline in Treg induction was observed in middle-aged slices expressing the less abundant RIP-OVA^hi^ TRA, in which mature Tregs were almost undetectable, representing less than 1% of the OT-II CD4SP thymocytes (**Figure 4d**). Together, these results demonstrate that the middle-aged thymic environment is impaired in its capacity to support Treg generation to TRAs but maintains the capacity to support Treg induction to abundant ubiquitous self-antigens.

### Aging associated changes in central tolerance of polyclonal thymocytes

To determine if the decline in central tolerance we observed with antigen-specific models is also evident in polyclonal thymocytes by 12MO of age, we quantified negative selection and Treg induction in mice from 1MO to 12MO of age. We assessed the frequency of post-positive selection thymocyte subsets undergoing apoptosis, as identified by intracellular cleaved caspase 3 (CCasp3). CD4^+^CD8^+^ double positive (DP) thymocytes were subdivided into early post-positive selection CD3^lo^CD69^+^ cells and later CD3^+^CD69^+^ cells. CD4SP subsets were divided into semimature (SM), mature 1 (M1), and mature (M2) cells, and CD8SPs were subset into M1 and M2 stages based on expression of CD69 and MHC-I (Xing et al. 2016) (**Figure 5a**). Each subset was gated for expression of CD5 and CD3 to ensure quantification of cells that had received a TCR signal, and the frequency of Ccasp3+ cells was then quantified. In general, the less mature DP CD3^lo^CD69^+^cells and CD4SM subsets underwent higher rates of negative selection than more mature SP subsets. Although there was a decline in the frequency of CD8SP M1 cells undergoing negative selection at 6MO of age, we did not observe a decline in the frequency of any polyclonal subsets undergoing negative selection by 12MO of age (**Figure 5b**). Thus, overall rates of negative selection are relatively constant in the thymus from 1MO through 12MO of age, likely reflecting ongoing negative selection to abundant ubiquitous self-antigens, in keeping with our findings that OT-I and OT-II thymocytes were deleted fairly efficiently to high avidity ubiquitous self-antigens.

**Figure 5.**
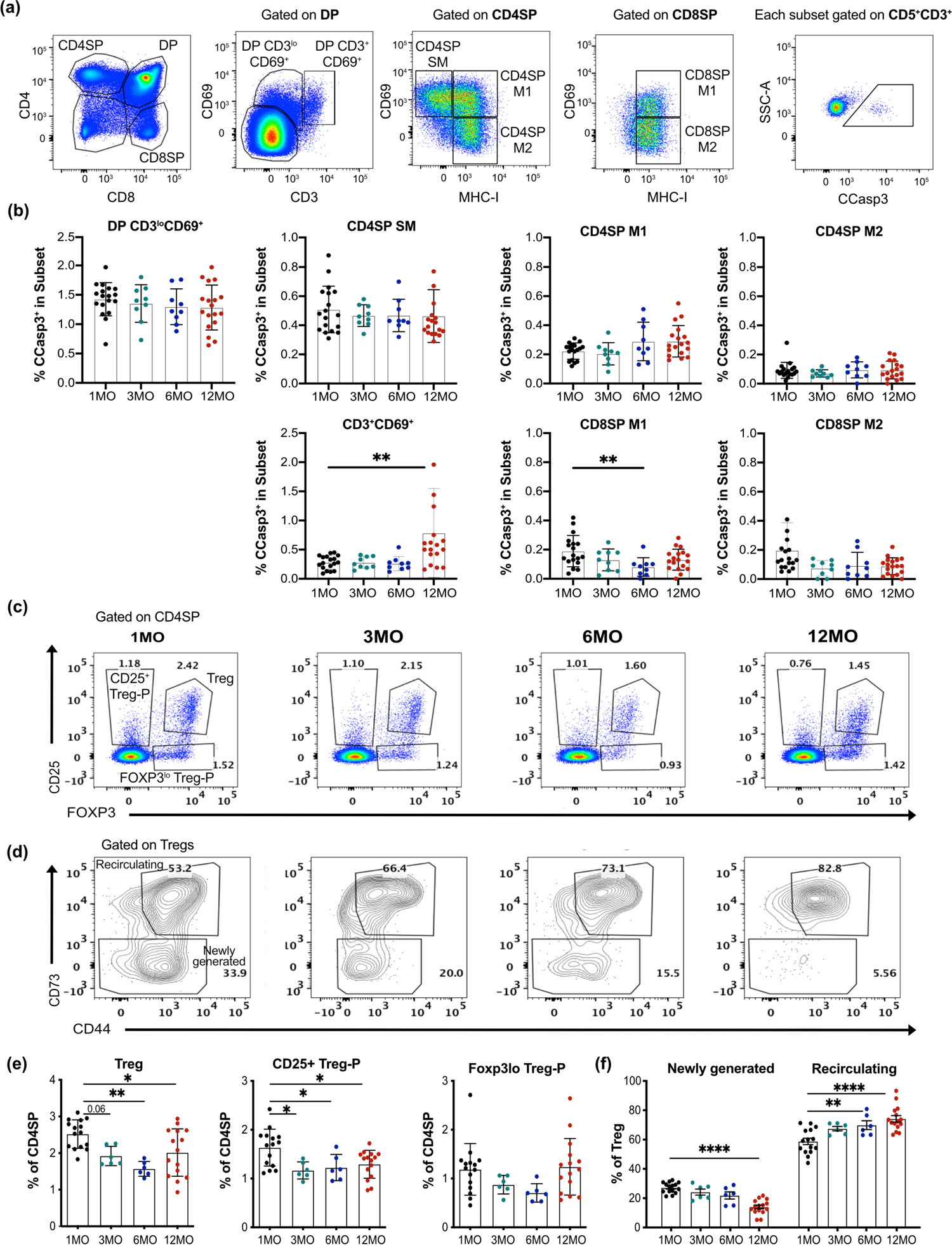
Aging associated changes in central tolerance of polyclonal thymocytes. **(a)** Flow cytometry gating strategy for identification of thymocyte subsets undergoing negative selection in the thymus. Postpositive selection DP thymocytes were subdivided into DP CD3^lo^CD69^+^ and DP CD3^+^CD69^+^ stages, while CD4SP and CD8SP cells were divided into semimature (SM), mature1 (M1) and mature2 (M2) stages, as indicated. Cells were subsequently gated on CD5^+^ CD3^+^ cells to restrict analysis to thymocytes that had undergone TCR signaling, and cleaved caspase 3 (CCasp3) expression identified cells undergoing clonal deletion in each subset. **(b)** Quantification of the frequency of CCasp3+ cells in each thymocyte subsets from mice at 1, 3, 6 and 12 MO of age. **(c)** Representative flow cytometry plots of Treg-P and Tregs at 1, 3, 6 and 12 MO of age. **(d)** Representative plots of CD4^+^CD25^+^Foxp3^+^ Tregs gated to distinguish newly generated (CD73^-^) from recirculating cells (CD73^+^) in thymuses from mice at 1, 3, 6 and 12 MO of age. **(e)** Quantification of Tregs, CD25^+^ Treg-Ps, and FOXP3^lo^ Treg-Ps expressed as a frequency of CD4SPs. **(h)** Quantification of newly generated versus recirculating Tregs expressed as a frequency of mature Tregs. (b, e-f) Plots show mean ± SEM of nine to fifteen thymuses per age. Symbols represent individual thymuses. Analyzed by one-way ANOVA with Tukey’s test for multiple comparisons, *p*-values: * < 0.05, ** < 0.01, *** < 0.001, **** < 0.0001.

We also tested if there was a decline in the generation of polyclonal Tregs and Treg-P by 12MO of age (**Figure 5c**). Previous studies indicated that *de novo* Treg induction in an aging thymus is impaired due to an increased number of peripheral Treg that recirculate into the thymus where they outcompete newly differentiating Treg for limited, local IL-2, which is required for Foxp3 upregulation (Hemmers et al. 2019; Thiault et al. 2015). To quantify de novo polyclonal Treg generation with age, we distinguished newly generated from recirculating Treg based on expression of CD73 (Owen et al. 2019) (**Figure 5d**). The overall frequency of Treg within the CD4SP compartment diminishes significantly by 6MO of age and remains low at 12MO (**Figure 5e**). Within the thymic Treg compartment, the frequency of newly generated cells steadily declines over 12MO of age, with a concomitant increase in recirculating Treg (**Figures 5d,f**), consistent with previous studies (Thiault et al. 2015). These findings indicate that the age-associated decrease in generation of new Treg can be detected in the polyclonal repertoire, consistent with our observation that Treg induction was severely compromised for OT-II thymocytes responding to TRAs and was somewhat impaired for cells responding to a ubiquitous a self-antigen in a 12MO thymic environment (**Figure 4**).

The proportion of CD25^+^ Treg-P within the CD4SP compartment decreases significantly with age, with a trend towards diminished frequencies of Foxp3^lo^ Treg-P as well (**Figure 5e**). Because CD25 upregulation is induced on thymocytes encountering cognate-self antigens by TCR stimulation (Lio & Hsieh 2008), the aging-associated reduction in CD25^+^ Treg-P is consistent with an age-associated decrease in thymocyte access to self-antigens that promote Treg differentiation. The concept that self-antigen availability limits the induction of Treg in a middle-aged thymus is concordant with the finding that OT-II thymocytes generate Treg-P fairly efficiently in a 12MO thymic environment in response to abundant ubiquitous self-antigens, but not to lower abundance endogenous TRAs (**Figure 4**).

### The composition of TEC and HAPC compartments is significantly altered in a middle-aged thymus

Given the reduced capacity of the middle-aged thymus to support efficient negative selection and Treg induction, particularly against self-antigens of moderate avidities, we tested whether aging altered the cellular composition of TECs and HAPCs, the major APC subsets critical for promoting central tolerance to diverse self-antigens. TECs, B cells, pDCs, cDCs, and macrophages were quantified by flow cytometric analysis of enzymatically digested thymuses at 1, 3, 6, and 12MO of age. cDCs were divided into cDC1 and cDC2 subsets based on expression of XCR1 and SIRP*α*, respectively, while mTECs were divided from cTECs based on mTEC expression of UEA-1. Both TEC and cDC subsets were subdivided by low and high levels of MHC-II expression, and CD80^+^MHCII^hi^ mTECs were further subdivided based on expression of AIRE (representative TEC and HAPC gating in **Supplementary Figure 3**). The number of TEC^lo^ cells steadily increased from 1MO to 6MO, with a slight reduction in cTEC^hi^ numbers by middle age (**Figure 6a**). Increased TEC^lo^ numbers were reflected in their increased representation within the overall cTEC population, with a commensurate decrease in cTEC^hi^ frequencies by middle-age (**Figure 6a**). All mTEC subsets declined numerically at 6 and 12MO of age, consistent with the overall decline in thymus cellularity, but only the AIRE^+^ mTEC^hi^ subset was reduced in frequency within the mTEC compartment, while mTEC^lo^ cells increased proportionally (**Figure 6a**). Thus, we find that by middle-age thymic involution is associated with a shift in TEC composition towards TEC^lo^ and mTEC^lo^ subsets, with a substantial decrease in the frequency of AIRE^+^ mTECs, the TEC subset that expresses diverse TRAs.

**Figure 6.**
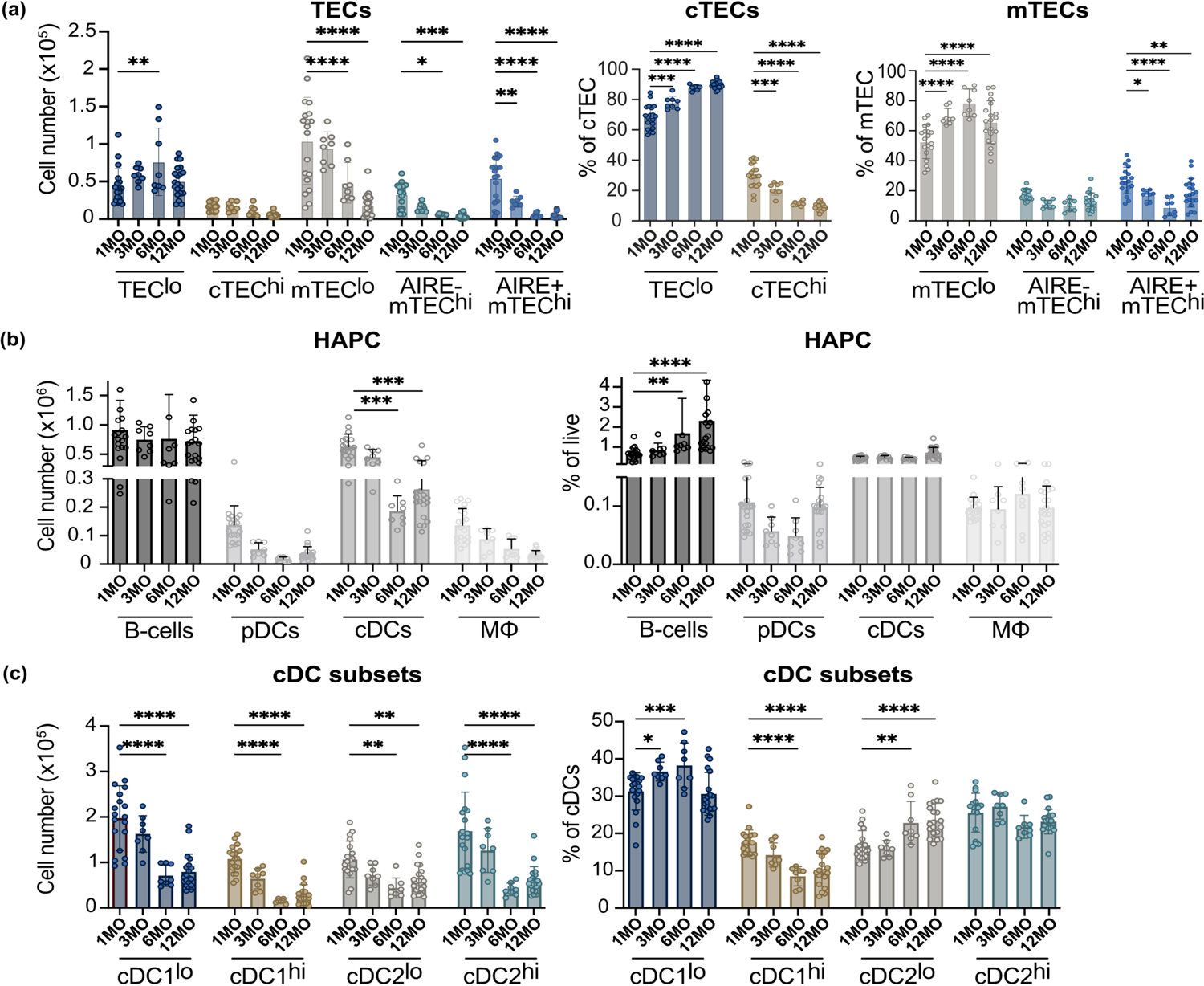
The composition of TEC and HAPC compartments is significantly altered in a middle-aged thymus. **(a)** Total cellularity and frequency of TEC subsets in 1, 3, 6 and 12MO thymuses were quantified by flow cytometry. **(b)** The total number of HAPC subsets and percent of live cells was quantified in 1, 3, 6 and 12MO thymus. TECs, B cells, pDC, cDCs and macrophages (m*Φ*) were gated as shown in **Supplementary** Figure 3. **(c)** The total number of thymic cDC subsets and percentage of live cells were quantified at 1, 3, 6 and 12MO of age. Plots show mean ± SEM of 8-20 thymuses per age. Data are compiled from 6 experiments. Analyzed by *t*-test, *p*-values: * < 0.05, ** < 0.01, *** < 0.001, **** < 0.0001.

The HAPC compartment also undergoes significant composition changes by middle-age. The number of B cells does not decline by 12MO, resulting in an increased frequency of total thymocytes (**Figure 6b**), consistent with previous studies (Cepeda et al. 2018). There was a trend towards decreasing plasmacytoid dendritic cell (pDC) numbers and frequencies with age (**Figure 6b**). cDCs numbers decreased significantly by 6 and 12 MO of age (**Figure 6b**). The rate of cDC attrition corresponded to the overall decline in thymic cellularity during involution, as their frequency remained constant with age (**Figure 6b**). Similarly, macrophages decreased in numbers but not in frequency by middle-age (**Figure 6b**). Overall, broad HAPC subsets decreased at approximately the same rate as thymic involution through middle-age, with the exception of thymic B cells, which increased in frequency.

As distinct cDC subsets have been differentially implicated in presenting self-antigens to induce tolerance (Leventhal et al. 2016; Perry et al. 2014; Perry et al. 2018; Oh et al. 2017; Ardouin et al. 2016), age-associated changes in cDC subsets were further evaluated. Within XCR1^+^ cDC1s, the number of both cDC1^lo^ and cDC1^hi^ subsets decreased significantly by 12MO of age. Interestingly, however, the frequency of cDC1^hi^ cells decreased within the cDC compartment, while that of cDC1^lo^ cells increased with age (**Figure 6c**). cDC1^hi^ cells have been implicated in acquiring self-antigens from *Aire*^+^ mTECs to induce Treg selection (Perry et al. 2014). For cDC2, both cDC2^lo^ and cDC2^hi^ numbers decreased significantly with age (**Figure 6c**). However, cDC2^lo^ constituted an increasing proportion of total cDCs with age, while frequency of cDC2^hi^ remained constant (**Figure 6d**). cDC2s have also been implicated in presenting Aire-dependent antigens to induce Treg selection (Leventhal et al. 2016). Thus, similar to changes in the TEC compartment, the cDC compartment becomes enriched for MHCII^lo^ subsets for both cDC1 and cDC2 by middle age, consistent with reduced presentation of diverse self-antigens to induce thymocyte tolerance.

## Discussion

Here we identify specific defects in central tolerance that manifest in a middle-aged thymus. Following peak thymus size and T-cell output at 1MO of age in mice, thymic cellularity and T-cell export progressively decline (Srinivasan et al. 2021; Chinn et al. 2012). During age-associated thymic involution, TECs become less proliferative, their numbers diminish, and their cellular composition and organization change. The number of AIRE+ mTECs and expression of TRAs decline in a 12MO, middle-aged thymic environment (Lepletier et al. 2019; Baran-Gale et al. 2020; Bredenkamp et al. 2014; Venables et al. 2019; Griffith et al. 2012; Gray et al. 2006), suggesting central tolerance to TRAs could be particularly impaired in the involuting thymus, but this possibility had not been evaluated. In this study, we used naturally aged thymic slices to separate the age of thymus from that of thymocytes to directly test if a middle-aged thymic environment retains the capacity to support central tolerance to ubiquitous self-antigens and/or TRAs. We determine that if thymocytes encounter high avidity self-antigens presented on either MHC-I or MHC-II molecules, negative selection is largely preserved in a middle-aged thymus. However, the 12MO thymic environment does not induce efficient negative selection or Treg differentiation of thymocytes signaled by lower avidity self-antigens, regardless of whether they are presented ubiquitously throughout the thymus or in the medulla as TRAs. Thus, by middle-age, the thymus is selectively impaired in its capacity to induce central tolerance to moderate avidity self-antigens.

Thymocytes must efficiently enter the medulla to scan APCs that present the diverse self-antigens, including *Aire*-dependent TRAs, that establish broad self-tolerance to autoantigens throughout the body. After positive selection, thymocytes upregulate the chemokine receptors CCR7 and CCR4, which promote their accumulation within the medulla where the corresponding chemokine ligands are expressed (Hu et al. 2015; Ehrlich et al. 2009; Ueno et al. 2004). Thus, we considered the possibility that as the thymus ages, and TEC organization deteriorates, thymocyte accumulation in the medulla could be compromised. However, the middle-aged thymic environment supported robust accumulation of CD4SP thymocytes in the medulla, comparable to a young thymus. Interestingly, CCL21, which is the CCR7 ligand responsible for attracting thymocytes into the medulla (Kozai et al. 2017), is expressed at higher levels in the medulla of a middle-aged relative to a young thymus. Also, post-positive selection 12MO thymocytes express comparable levels of CCR4 and CCR7 to 1MO thymocytes. Thus, our data indicate that impaired central tolerance in the middle-aged thymus is not due to reduced entry or accumulation of single-positive thymocytes in the medulla.

Given that thymocyte medullary residence is limited to about 5 days (McCaughtry et al. 2007), and many self-antigens are displayed in a sparse mosaic in the thymus (Sansom et al. 2014; Meredith et al. 2015; Derbinski et al. 2008; Derbinski et al. 2005; Baran-Gale et al. 2020), thymocytes need to migrate rapidly within the medulla to encounter sufficient APCs to ensure complete tolerance. We find that the velocity of CD4SP migration is significantly reduced in a middle-aged 12MO versus a young 1MO thymic environment, regardless of thymocyte age. This reduction in migratory speed induced by the 12MO thymic environment could limit the number of APCs encountered, diminishing the efficiency of central tolerance. Future studies are needed to determine why thymocyte velocities are reduced in a 12MO thymus.

Because expression of TRAs declines with age, and our data along with previous studies show that both the absolute number of AIRE+ mTECs and their frequency within the mTEC compartment are reduced (Griffith et al. 2012; Lepletier et al. 2019; Baran-Gale et al. 2020; Bredenkamp et al. 2014), we hypothesized that an age-associated deficiency in negative selection would be most appreciable for TRAs. Our data are somewhat consistent with this conclusion in that the middle-aged thymic environment largely supported negative selection of OT-I CD8SP thymocytes to exogenously administered ubiquitous SIINFEKLp, while the efficiency of negative selection to the RIP-OVA^hi^ TRA was significantly reduced by about 30%. However, ubiquitous versus *Aire*-dependent tissue restricted expression of a self-antigen was not sufficient to predict whether tolerance induction would be impaired in a middle-aged thymic environment. For example, negative selection of OT-I CD8SP thymocytes to the RIP-mOVA TRA was intact in 12MO thymic slices, while negative selection of OT-II CD4SP thymocytes to ubiquitous OVAp was impaired.

A better predictor of whether negative selection is impaired in a middle-aged thymic environment is the avidity of the selecting self-antigen for the TCR, which is impacted by both the amount of self-antigen encountered by thymocytes and the TCR-binding affinity of the self-antigen. We first evaluated negative selection using altered peptide ligands for the OT-I TCR that are reported to be just below (T4), just above (Q4R7), or well above (SIINFEKL) the TCR affinity threshold for clonal deletion (Daniels et al. 2006). OT-I negative selection in the middle-aged thymus was intact in response to high concentrations of all three self-antigens but was impaired in the presence of low concentrations of both high and low affinity peptides. Moreover, only the lowest concentration of the high affinity OT-I self-antigen revealed the deficit in negative selection in the middle-aged thymus, while this deficiency was apparent at relatively higher concentrations of lower affinity self-antigens. These findings show that the middle-aged thymic environment is less efficient at displaying ubiquitous self-antigens to induce negative selection, effectively raising the avidity threshold for negative selection by middle-age. The mechanisms underlying the deficit in negative selection remain to be determined, but could reflect changes in APC composition, such as the reduced frequency of MHCII^hi^ cDCs.

The middle-aged thymus was also deficient in inducing negative selection against lower avidity TRAs. Negative selection of OT-I thymocytes in 12MO thymic environments was intact against the higher affinity RIP-mOVA TRA, but was significantly impaired against the lower affinity variants RIP-mQ4R7 and RIP-mT4, as well as the high affinity RIP-OVA^hi^ TRA, which is expressed at lower levels than RIP-mOVA (Lancaster et al. 2019; Kurts et al. 1998). Taken together, these findings further support that negative selection to lower avidity self-antigens, even when expressed as TRAs, is impaired in the middle-aged thymus. Mechanisms underlying diminished negative selection to TRAs have yet to be determined, but could reflect a combination of the lower velocity of thymocytes, as discussed above, as well as the significant reduction in both the numbers and frequencies of AIRE^+^ mTECs that express and present TRAs (Aschenbrenner et al., 2007; Hinterberger et al., 2010; Lancaster et al., 2019) and the MHCII^hi^ cDC1s that have been implicated in acquiring antigens from Aire+ mTECs for display to thymocytes (Perry et al. 2014; Perry et al. 2018).

Although the decline in negative selection observed in the middle-aged thymus may seem modest, even a small number of self-reactive T cells in the periphery can induce autoimmunity (Bosch et al. 2017). Interestingly, when the low affinity T4p was expressed as a TRA or presented as a low-level ubiquitous self-antigen in 12MO thymic slices, we consistently recovered more OT-I CD8SPs than in control slices without antigen. T4p has been shown to induce negative selection at high concentrations and positive selection at lower doses (Daniels et al. 2006). Thus, by middle-age the thymus may induce positive selection of thymocytes specific for low avidity self-antigens, perhaps resulting in export of an elevated number of autoreactive T cells to the periphery. Importantly, OT-I CD8 T cells that escaped negative selection to such low avidity TRAs (RIP-mQ4R7 and RIP-mT4) potently induced autoimmune diabetes upon immunization with OVAp (Koehli et al. 2014). Future experiments will investigate if OT-I CD8SP that escape negative selection in the middle-aged thymus more potently prime such autoimmune responses. Furthermore, although we did not find evidence of impaired negative selection of polyclonal thymocytes at 12 MO of age, the frequency of CD8SP M1 cells undergoing negative selection declined at 6MO. Because negative selection in the middle-aged thymus is impaired selectively to moderate avidity self-antigens, intact negative selection to abundant high-avidity self-antigens could have masked the deficit in polyclonal cells. Altogether, these findings suggest that impaired negative selection in a middle-aged thymus could result in export of weakly autoreactive T cells that have the potential to induce autoimmunity, potentially contributing to the peak in new-onset autoimmunity in middle-age (Watad et al. 2017). Future studies will test this possibility.

The middle-aged thymic environment was particularly impaired in supporting generation of new Treg responsive to TRAs; however, Treg differentiated efficiently in response to ubiquitous antigens. The 12MO thymic environment supported efficient Treg generation against the ubiquitous self-antigen OVAp, which was surprising as previous studies suggested that new Treg development is limited by IL-2 availability, due to competition from recirculating peripheral Tregs (Weist et al. 2015; Thiault et al. 2015). However, despite the presence of more recirculating Treg in the 12MO thymic environment, sufficient IL-2 was present to induce robust Treg generation in the context of exogenously added OVAp. Tregs in the thymus can arise through two distinct progenitors, CD25+ Treg-P or FOXP3^lo^ Treg-P (Owen et al. 2019). In a two-step process, thymocytes first receive a TCR signal that upregulates the high-affinity IL-2 receptor α-chain (CD25), generating CD25^+^ Treg-P progenitors. Subsequent IL-2 signaling induces *Foxp3* expression and Treg differentiation (Lio & Hsieh 2008; Burchill et al. 2008).

Alternatively, thymic Tregs can arise via FoxP3^lo^ Treg-P, which differ from CD25+ Treg-P in their transcriptome, TCR repertoire, developmental kinetics, susceptibility to apoptosis, dependence on cytokines, and suppressive functions (Tai et al. 2013; Owen et al. 2019). Exogenous OVAp induced robust CD25 expression in 1MO and 12MO thymi, indicating that the middle-aged thymus can efficiently present ubiquitous self-antigens to MHCII-restricted thymocytes. If IL-2 were limiting in the 12MO thymus due to an increase in recirculating Treg, fewer CD25^+^ Treg-P would be expected to upregulate FOXP3 and differentiate into Treg. However, the 12MO environment generated a comparable number of Treg as the 1MO environment, despite increased numbers of recirculating Treg. In contrast, there was a significant reduction in the frequency of OT-II CD4SP that differentiated into FOXP3^lo^ Treg-P in the 12MO thymic environment in response to RIP-mOVA and RIP-OVA^hi^ TRAs. Interestingly, the RIP-mOVA TRA induced the most robust differentiation of Foxp3^lo^ Treg-P, with ∼17% of CD4SPs exhibiting this phenotype in the 1MO environment. Notably, the number and frequency of OT-II Tregs decline substantially in response to both model TRAs; mature Treg induction to RIP-OVA^hi^ in the middle-aged thymus environment was virtually extinguished. Altogether, these findings, in combination with the robust Treg generated in response to ubiquitous OVAp, are more consistent with a model in which the middle-aged thymus has a deficiency in the Treg niche that presents TRAs to CD4SP thymocytes. However, access to other cytokines that can contribute to Treg differentiation, like IL-4(Owen et al. 2019), or local concentrations of IL-2 could also be limiting. The reduced frequencies of both AIRE^+^ mTECs and MHCII^hi^ cDC1s in the middle-aged thymic environment, both of which express and/or present *Aire*-dependent TRAs to autoreactive thymocytes (Perry et al. 2014; Perry et al. 2018; Lancaster et al. 2019; Ardouin et al. 2016; Hinterberger et al. 2010; Aschenbrenner et al. 2007; Hubert et al. 2011; Gallegos & Bevan 2004), suggest that an altered APC compartment is a major contributor to the reduction in selection of new Tregs in the aging thymic environment. However, it remains to be resolved how age-associated changes in APC subsets impact negative selection and Treg induction to distinct self-antigens in the young versus aged thymus.

### Experimental Procedures

#### Mice

C57BL/6J (Jackson Laboratories and NIH/NIA), B6.SJL-Ptprc^a^Pepc^b^/BoyJ (CD45.1), C57BL/6-Tg(TcraTcrb)1100Mjb/J (OT-I)(Hogquist et al. 1994), B6.Cg-Tg(TcraTcrb)425Cbn/J (OT-II)(Barnden et al. 1998), C57BL/6-Tg(Ins2-TFRC/OVA)296Wehi/WehiJ (RIP-mOVA)(Kurts et al. 1996), RIP-OVA^hi^ (W. R. Heath, University of Melbourne, Melbourne, Australia)(Kurts et al. 1998), RIP-mT4 (E. Palmer, University of Basel, Basel, Switzerland), RIP-mQ4R7 (E. Palmer, University of Basel, Basel, Switzerland) and pCX-EGFP (I. Weissman, Stanford University, Stanford, CA)(Wright et al. 2001) strains were bred in-house. All strains were sourced from Jackson Laboratories, except as specified. Mouse maintenance and experimental procedures were carried out with approval from the Institutional Animal Care and Use Committee at the University of Texas at Austin. All strains were bred and maintained under specific pathogen-free conditions in the University of Texas at Austin animal facility.

#### Antibodies

Antibodies used in flow cytometry and immunofluorescence analyses are detailed in **Table 1**. For flow cytometry, 5-10 *×* 10^6^ cells were immunostained in 100 μL of PBS + 2% bovine calf serum (BCS) with fluorochrome-conjugated antibodies. Unless otherwise specified, cells were incubated with fluorochrome-conjugated antibodies for 20 min on ice, washed twice in PBS + 2% BCS, incubated with streptavidin for 20’ on ice, washed and resuspended in 10 μg mL^-1^ propidium iodide (PI) to determine viability.

**Table 1.**
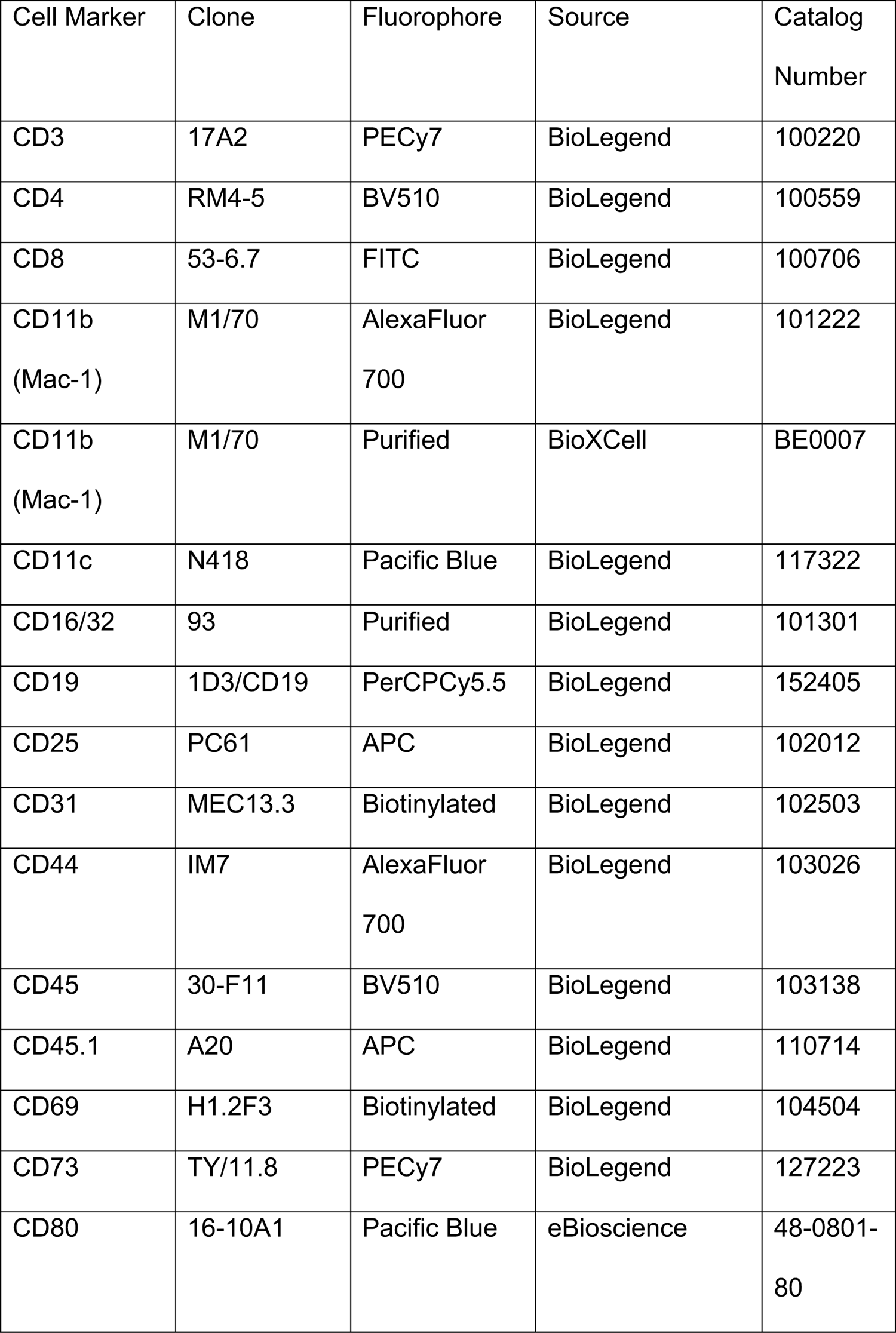
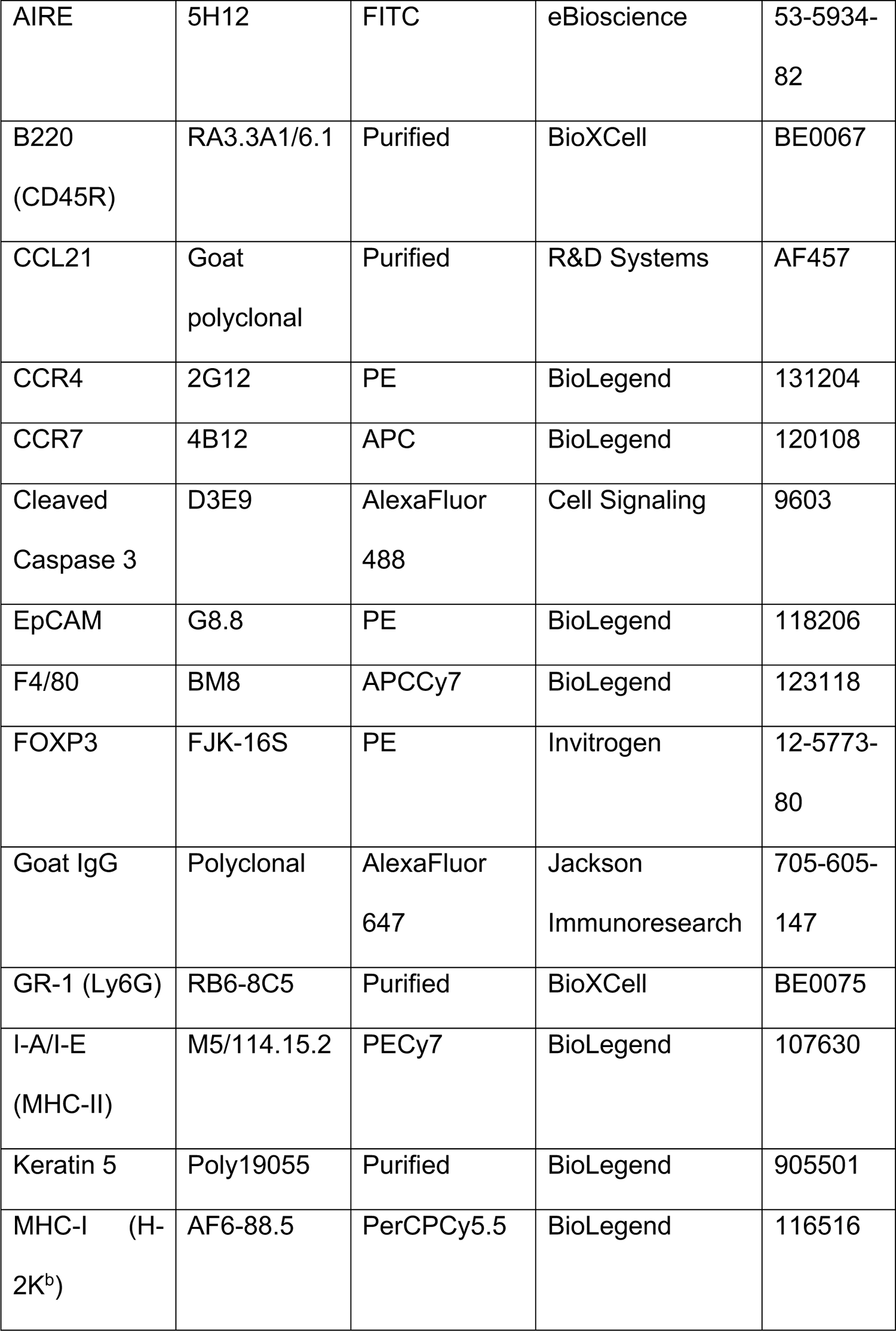
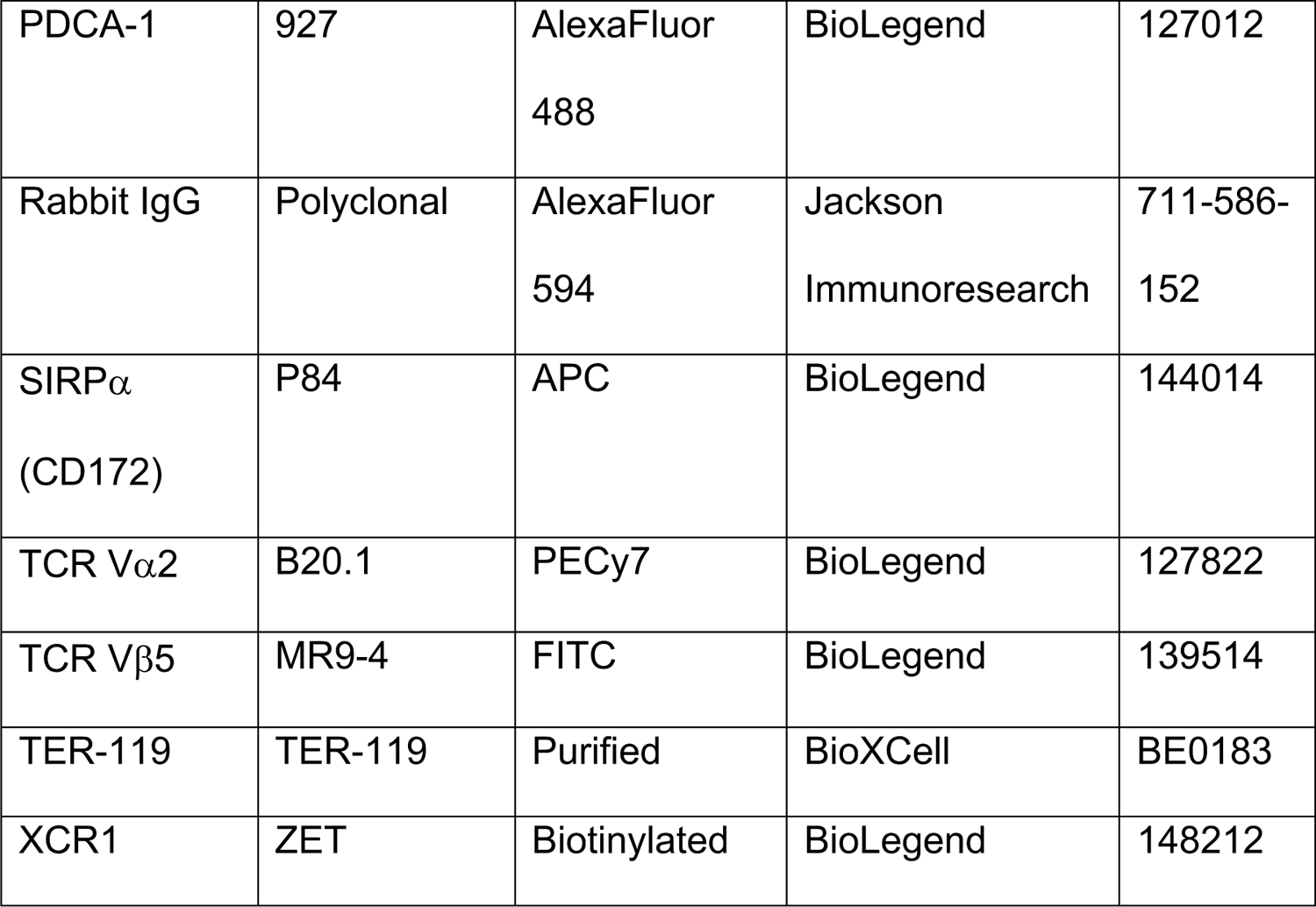

| Cell Marker | Clone | Fluorophore | Source | Catalog Number |
| --- | --- | --- | --- | --- |
| CD3 | 17A2 | PECy7 | BioLegend | 100220 |
| CD4 | RM4-5 | BV510 | BioLegend | 100559 |
| CD8 | 53-6.7 | FITC | BioLegend | 100706 |
| CD11b<br>(Mac-1) | M1/70 | AlexaFluor<br>700 | BioLegend | 101222 |
| CD11b<br>(Mac-1) | M1/70 | Purified | BioXCell | BE0007 |
| CD11c | N418 | Pacific Blue | BioLegend | 117322 |
| CD16/32 | 93 | Purified | BioLegend | 101301 |
| CD19 | 1D3/CD19 | PerCPCy5.5 | BioLegend | 152405 |
| CD25 | PC61 | APC | BioLegend | 102012 |
| CD31 | MEC13.3 | Biotinylated | BioLegend | 102503 |
| CD44 | IM7 | AlexaFluor<br>700 | BioLegend | 103026 |
| CD45 | 30-F11 | BV510 | BioLegend | 103138 |
| CD45.1 | A20 | APC | BioLegend | 110714 |
| CD69 | H1.2F3 | Biotinylated | BioLegend | 104504 |
| CD73 | TY/11.8 | PECy7 | BioLegend | 127223 |
| CD80 | 16-10A1 | Pacific Blue | eBioscience | 48-0801-80 |
| AIRE | 5H12 | FITC | eBioscience | 53-5934-82 |
| B220<br>(CD45R) | RA3.3A1/6.1 | Purified | BioXCell | BE0067 |
| CCL21 | Goat<br>polyclonal | Purified | R&D Systems | AF457 |
| CCR4 | 2G12 | PE | BioLegend | 131204 |
| CCR7 | 4B12 | APC | BioLegend | 120108 |
| Cleaved<br>Caspase 3 | D3E9 | AlexaFluor<br>488 | Cell Signaling | 9603 |
| EpCAM | G8.8 | PE | BioLegend | 118206 |
| F4/80 | BM8 | APCCy7 | BioLegend | 123118 |
| FOXP3 | FJK-16S | PE | Invitrogen | 12-5773-80 |
| Goat IgG | Polyclonal | AlexaFluor<br>647 | Jackson<br>Immunoresearch | 705-605-147 |
| GR-1 (Ly6G) | RB6-8C5 | Purified | BioXCell | BE0075 |
| I-A/I-E<br>(MHC-II) | M5/114.15.2 | PECy7 | BioLegend | 107630 |
| Keratin 5 | Poly19055 | Purified | BioLegend | 905501 |
| MHC-I (H-2K<sup>b</sup>) | AF6-88.5 | PerCPCy5.5 | BioLegend | 116516 |
| PDCA-1 | 927 | AlexaFluor<br>488 | BioLegend | 127012 |
| Rabbit IgG | Polyclonal | AlexaFluor<br>594 | Jackson<br>Immunoresearch | 711-586-152 |
| SIRPα<br>(CD172) | P84 | APC | BioLegend | 144014 |
| TCR Vα2 | B20.1 | PECy7 | BioLegend | 127822 |
| TCR Vβ5 | MR9-4 | FITC | BioLegend | 139514 |
| TER-119 | TER-119 | Purified | BioXCell | BE0183 |
| XCR1 | ZET | Biotinylated | BioLegend | 148212 |

#### Purification of thymocytes for live-cell microscopy

CD4SP cells were enriched by incubating 2 *×* 10^8^ cells mL^-1^ with antibodies against CD8 (96 μg mL^-1^) and CD11b, GR-1, TER-119, and CD25 (5 μg mL^-1^ each) for 30 min on ice in PBS + 2% BCS, followed by immunomagnetic depletion using sheep anti-rat IgG magnetic DynaBeads (Life Technologies at a 2:1 cell:bead ratio. Magnetic depletion was repeated with half the number of beads to improve enrichments. Purity of isolated CD4SP was determined by flow cytometry using the following fluorochrome-conjugated antibodies: anti-CD3, -CD4, -CD8, and -CD69, followed by Streptavidin Qdot605 (Thermo Fisher Scientific, 1:800 dilution). Cells were washed and resuspended in 10 μg mL^-1^ propidium iodide (PI) to determine viability. Samples were analyzed on an LSR Fortessa flow cytometer (BD Biosciences), and data were analyzed using FlowJo (v.10, TreeStar). For each slice, 10^6^ isolated cells were stained with either 2 μM CMTPX CellTracker Red or 2 μM Indo1AM (both from Life Technologies) for 30 min at 37°C in 1.5 mL of DRPMI medium (RPMI 1640 without L-glutamine, phenol red, and sodium bicarbonate; Cellgro) supplemented with 0.2 g L^-1^ sodium bicarbonate and 20 mM HEPES. Cells were washed and incubated in 1.5 mL complete RPMI medium (RPMI 1640 with 2 mM L-glutamine, 50 U mL^-1^ penicillin, 50 mg mL^-1^ streptomycin, and 10% (v/v) fetal bovine serum) for 30 min to destain. Cells were washed, combined so that 10^6^ CellTracker Red-labeled cells and 10^6^ Indo1AM-labeled cells were mixed into each tube, and washed again with complete RPMI medium. Thymocytes were concentrated into 20-μL complete RPMI medium and carefully pipetted onto the surface of each thymic slice before incubation at 37°C 5% CO_2_ to allow migration of thymocytes into the thymic slice. To avoid color channel bias, 1MO and 12MO thymocyte fluorophores were swapped in different experiments.

#### Thymic slice preparation

For 2PM imaging, slices were generated from pCX-EGFP thymi. For negative selection assays, slices were generated from C57BL/6, RIP-mOVA, RIP-OVAhi, RIP-mT4, or RIP-mQ4R7 thymi. Dissected thymi were embedded in 4% (w/v) NuSieve GTG low-melting-temperature agarose (Lonza) in PBS at 37°C. The solidified agarose block was sectioned into 400-μm-thick slices using a VT 1000 S Microtome (Leica) in a bath of ice-cold PBS, with vibratome frequency set to 70 Hz, speed to 0.20 mm s^-1^, and amplitude to 0.6 mm. Slices were collected in DRPMI + 10% bovine calf serum on ice before transfer to 0.4-μm tissue culture inserts (Millipore) in 35-mm Petri dishes containing 1 mL of complete RPMI medium, with or without added peptides.

#### Two-photon fluorescence microscopy

After incubation for *≥*1 h, pCX-EGFP thymic slices were transferred and secured in an imaging chamber (Harvard Apparatus) on the microscope stage. Perfusion medium, consisting of DRPMI supplemented with 2 g L^-1^ sodium bicarbonate, 5 mM HEPES, and 1.25 mM calcium chloride, was gravity fed to the stage inlet through a 300-mL IV set, and circulated through the imaging chamber at a flow rate at ∼100 mL h^-1^, or ∼1 drop per second. The perfusion medium was aerated with 95% oxygen and 5% carbon dioxide and maintained at 37°C with a heated microscope stage and inline perfusion heater. Images were acquired every 15 s, through a depth of 40 μm, at 5-μm intervals for durations of 15 min, using an Ultima IV microscope (Bruker) with a 20*×*water immersion objective (NA 1.0) and PrairieView software (v.5.4, Bruker). The sample was illuminated with two MaiTai titanium:sapphire lasers (Newport) tuned to 750 nm (for Indo1AM) and 900 nm (for CMTPX and EGFP). Emitted light was passed through 473/24, 525/50, and 605/70 band-pass filters (Chroma) to separate GaAsP detectors for detection of Indo1 (blue), EGFP, and CMTPX (red) fluorescence, respectively.

Migratory paths for thymocytes were tracked, and mean cell velocity and path straightness calculated using Imaris (v9, Bitplane). The enrichment of thymocytes in the medulla was determined at the first time point for each dataset by measuring the number of thymocytes in manually demarcated cortical and medullary regions.

#### Negative selection assays in thymic slices

10^6^ OT-I or OT-II thymocytes and 10^6^ CD45.1 thymocytes per slice, along with the input control, were stained in 5 mL of DRPMI medium supplemented with 0.2 g L^-1^ sodium bicarbonate and 20 mM HEPES, with 5 μM CMF2HC CellTracker Blue (Life Technologies). Cells were washed and resuspended in 5 mL of complete RPMI medium for 30 min to destain, and then washed twice before application onto thymic slices generated from C57BL/6J, RIP-mOVA, RIP-OVA^hi^, RIP-mT4, or RIP-mQ4R7 mice. Slices were incubated in 37°C/5% CO_2_ on tissue culture inserts in Petri dishes containing 1 mL of complete RPMI, with or without OVAp (OVA_257-264_ for OT-I, New England Peptide; or OVA_323-339_ for OT-II, GenScript), T4p (Anaspec) or Q4R7p (GenScript) peptides for specified durations.

For analysis, slices were gently washed twice by submerging in PBS and manually disrupted to obtain single-cell suspensions. Input thymocytes and slice samples were stained with the following fluorophore-conjugated antibodies: anti-CD3, -CD4, -CD8, -CD25, -TCR V*α*2, -TCR V*β*5, and -CD45.1 (Table 1). Samples were washed and resuspended in 10 μg mL^-1^ PI for viability. For Treg induction experiments, samples were stained with fluorophore-conjugated antibodies against surface markers and Zombie Red viability dye (BioLegend) as described above, and fixed and permeabilized using the FOXP3/Transcription Factor Fix Perm kit (Tonbo Biosciences) per manufacturer instructions. Intracellular FOXP3 was stained by fluorophore-conjugated antibody for 20 min on ice, washed, and resuspended in PBS. After staining, 5 *×* 10^4^ polystyrene beads were added to the tubes for cell quantification and flow cytometric analysis was carried out. Cell subsets were quantified and normalized for variable slice entry based on the ratio of control CD45.1^+^ cells in each slice to the comparable CD45.1^+^ cells in the input sample. Triplicate slices of each condition were analyzed in each experiment. Data were normalized to the average number of cells in OVA-slices in the same experiment.

#### Flow-cytometric analyses of thymic stroma

Dissected thymi were cut into small fragments and enzymatically digested in 2-mL PBS with 2.5 mg/mL Liberase (Roche) with 120 μL DNase I solution [10,000 U, Roche, in 10 mL 50% glycerol: 50% DNase buffer (40 mM Tris-HCl +100 mM NaCl + 200 μg mL^-1^ bovine serum albumin)] for 12 min at 37°C, gently swirling halfway through incubation. The supernatant was transferred into 35 mL of PBS + 2% bovine calf serum and 5 mM EDTA at 4°C, with digestion of the remaining tissue fragments repeated twice to completely dissociate the tissue. The cells were spun down and filtered through a 70-μm nylon mesh to achieve a single-cell suspension.

For TEC analyses, cells were stained with 1 μg mL^-1^ biotinylated *Ulex europaeus* agglutinin I (UEA-1; Vector Laboratories) and the following fluorophore-conjugated antibodies: anti-CD11c, -CD45, -CD80, -EpCAM, and -I-A/I-E, and Zombie Red viability dye. Secondary staining was conducted with streptavidin conjugated Qdot 605 (1:400, Invitrogen) on ice for 20 min, followed by washing and fixation and permeabilization with the FOXP3/Transcription Factor Fix Perm kit. Intracellular AIRE was stained by fluorophore-conjugated antibody for 30 min on ice, washed, and resuspended in PBS. Quantification of TEC subsets was conducted by flow cytometry.

For HAPC analyses, cells were stained with the following fluorophore-conjugated antibodies: anti-CD11b, -CD11c, -CD19, -F4/80, -I-A/I-E, -PDCA-1, -SIRP*α* -XCR1. After washing, secondary staining was conducted by incubating with streptavidin-conjugated Qdot 605 (1:400, Invitrogen) on ice for 20 min. Samples were washed and resuspended in 10 μg mL^-1^ PI for viability. Quantification of HAPC subsets was conducted by flow cytometry.

### Flow cytometric analyses of thymocytes

Single-cell suspensions of thymocytes were obtained by manually dissociating dissected thymi and filtering cells through 40-μm cell strainers. Samples were immunostained with the following fluorophore-conjugated antibodies: anti-CD3, -CD4, -CD8, -CD69, -MHC-I, -CCR4, -CCR7 for 30 min at 37°C, prior to washing and resuspending the cells in 10 μg mL^-1^ PI for viability. To quantify negative selection within the thymus, thymocytes were immunostained with the following fluorophore-conjugated antibodies and dye: anti-CD3, -CD4, -CD5, -CD8, -CD69, -MHC-I and Zombie Red Viability dye (1:1000, BioLegend) for 30 min on ice. Cells were washed, then fixed and permeabilized using the Cytofix/Cytoperm kit (BD Biosciences BDB554714) per manufacturer instructions. Intracellular cleaved caspase 3 was stained with fluorophore-conjugated anti-cleaved caspase 3 antibody for 30 min on ice, washed, and resuspended in PBS for flow acquisition. Quantification of thymocyte subsets was conducted by flow cytometry.

To analyze polyclonal Tregs, single-cell thymocyte suspension were immunostained with the following fluorophore-conjugated antibodies: anti-CD3, -CD4, -CD8, -CD25, -CD44, -CD73 and Zombie Red viability dye (1:1000 dilution, BioLegend) at 4°C, and then fixed and permeabilized using the FOXP3/Transcription Factor Fix Perm kit (Tonbo Biosciences) per manufacturer instructions. Intracellular FOXP3 was immunostained with a fluorophore-conjugated antibody for 20 min on ice, and cells were washed and resuspended in PBS. Quantification of thymocyte subsets was conducted by flow cytometry.

### Immunofluorescence analysis of thymic cryosections

Thymuses were embedded in Tissue-Tek OCT medium and frozen using a mixture of dry ice and isopentane. 7-μm cryosections were generated with CryoStar NX50 cryostat (ThermoFisher) and stored at −80°C. Prior to immunostaining, sections were fixed in 100% acetone at −20°C for 20 minutes, washed with PBS + 0.1% Tween-20, and blocked with a solution of 10mM Tris HCl, 150mM NaCl and 0.5% blocking reagent (Perkin Elmer TSA Biotin System kit component). Fc receptor blocking was carried out with a 30-min room temperature stain using 10 µg mL^-1^ *α*-CD16/32. The sections were then incubated overnight at 4 °C with 10 µg mL^-1^ *α*-CCL21, 5 µg mL^-1^ *α*-CD31-biotin, and 1.66 µg mL^-1^ *α*-Keratin 5. Following the overnight stain, the sections were incubated for 1 h at room temperature with fluorophore-conjugated secondary antibodies and streptavidin-conjugated Alexa Fluor 488 (Life Technologies, S11223). 4’,6-diamidino-2-phenylindole (DAPI; Life technologies) was used at 0.125 µg mL^-1^ in PBS to stain nuclei.

## Statistics

All statistical analyses were performed using Prism (GraphPad) with the corresponding test and multiple-test corrections listed in the Figure Legends.

## Acknowledgements

We thank Janko Nikolich-Zugich, Nancy Manley, Marcel van den Brink, Jarrod Dudakov, Gregory Sempowski, Laura Hale, Bonnie LaFleur, and Jen Uhrlaub for constructive feedback on experiments and manuscript preparation. This research was supported by a grant from the National Institutes of Health, P01AG052359, to L.I.R.E. and E.R.R.

## Author Contributions

J.L., D.K.-C., and L.E. designed the experiments and wrote the manuscript; J.L., D.K.-C., J.S., and Y.L. performed experiments and analyzed data; H.S. and S.N. performed experiments; E.R. and L.E. edited the manuscript.

## Data Availability Statement

The data used to support the findings of this study are included within the article. Imaging and other data that support the findings of this study are available from the corresponding author upon request.

## Supporting Information Listing

Supplementary Figure, related to Figure 1

Supplementary Figure 2, related to Figure 3

Supplementary Figure 3, related to Figure 6

Supplementary Movie 1, related to Figure 1

Supplementary Movie 2, related to Figure 1

Supplementary Methods

