## Supplementary Data for "Central tolerance is impaired in the middle-aged thymic environment"

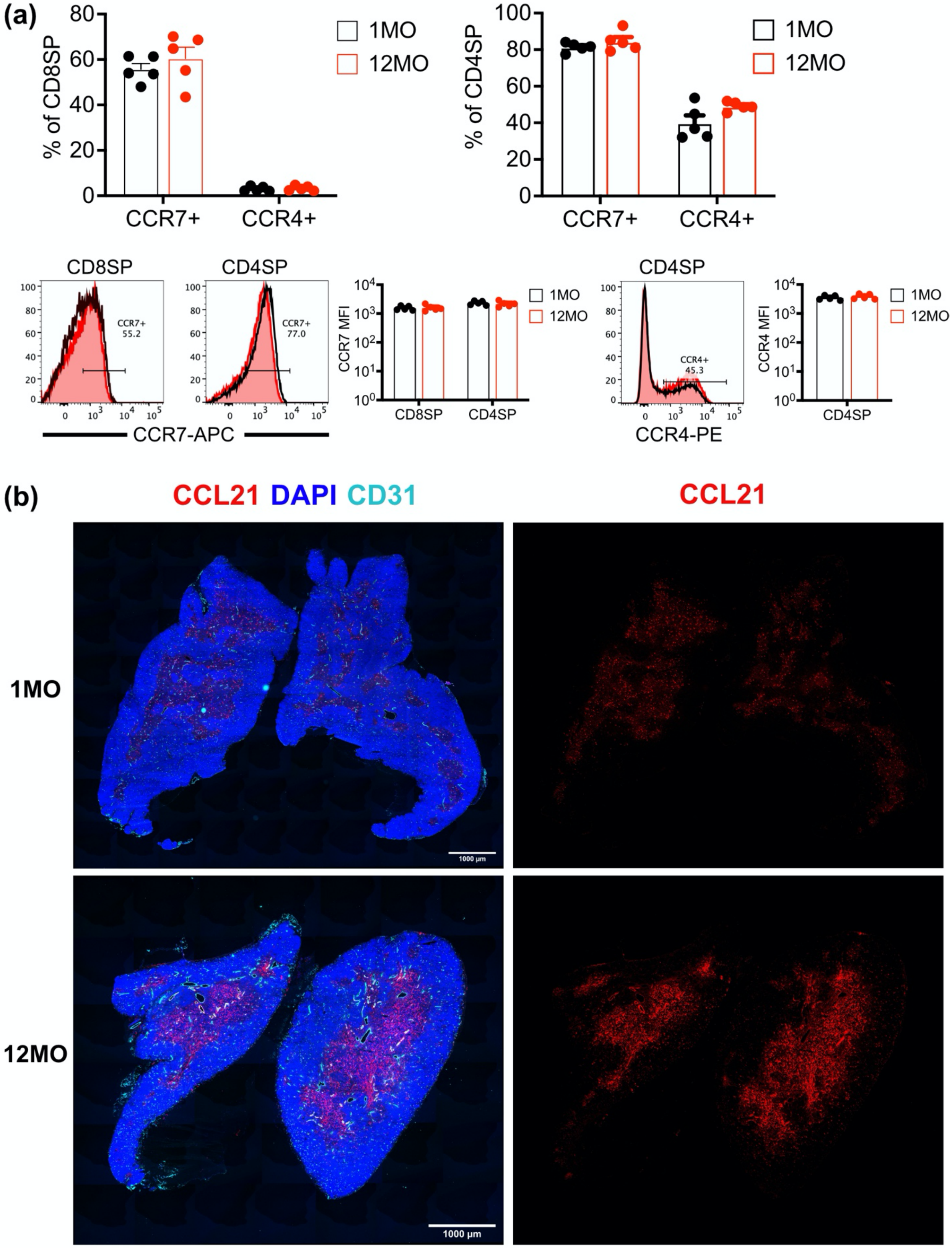

(a)
% of CD8SP
CCR7+
CCR4+
1MO
12MO
% of CD4SP
CD8SP
CD4SP
CCR7+
55.2
CCR7+
77.0
CCR7-APC
CCR7 MFI
CCR4+
45.3
CCR4-PE
CCR4 MFI
(b)
CCL21 DAPI CD31
CCL21
1MO
12MO
1000 μm

**Supplementary Figure 1, related to Figure 1. Expression of CCR4, CCR7, and CCL21 do not decline in 12MO versus 1MO thymuses. (a)** The proportion of CD8SP and CD4SP thymocytes that express the chemokine receptors CCR7 and CCR4 in 1MO (black) and 12MO (red) thymuses were evaluated by flow cytometry (top row). Histograms of CCR7 and CCR4 expression on CD8SP and/or CD4SP populations are shown, with gates delineating receptor-expressing cells; MFIs for receptor-positive cells were quantified(bottom row). The threshold for CCR7 expression was set based on lack of expression in $CD4^{-}CD8^{-}$ thymocytes. Each data point represents a mouse, with bars showing the means + SEM. **(b)** Immunofluorescence images of 1MO and 12MO thymus sections, immunostained for CCL21 (red) and CD31 (cyan), with a DAPI nuclear counterstain (blue). Staining and display parameters were kept constant to reveal differences in expression levels of CCL21 in 1 vs 12 mo thymi. Scale bar is 1000 μm.

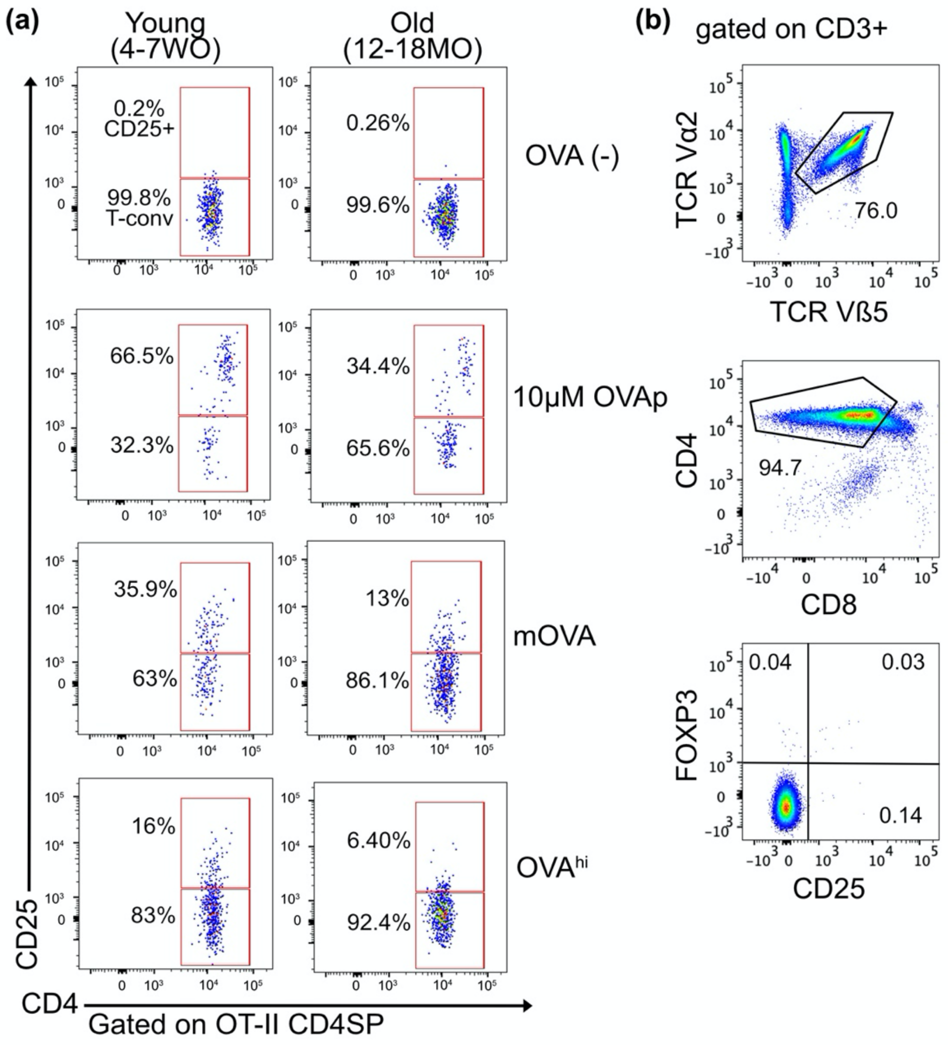


**Supplementary Figure 2, related to Figure 3. Deletion and Treg induction of OT-II CD4SPs against exogenous and endogenous self-antigens. (a)** Representative flow cytometry plots gated to show OT-II CD4SP CD25$^{-}$ and OT-II CD4SP CD25+ cells recovered from OVA(-), RIP-mOVA (mOVA) and RIP-OVA$^{hi}$ (OVA$^{hi}$) live thymic tissue slices from young vs. aged mice, at 48 h post-incubation. 10 μM of OVAp$_{323\text{-}339}$ (OVAp) was added as a positive control for OT-II negative selection. **(b)** Sequential gating scheme for flow cytometric analysis of OT-II thymocytes shows lack of Treg and Treg-P in input OT-II thymocytes used in Treg generation assays.

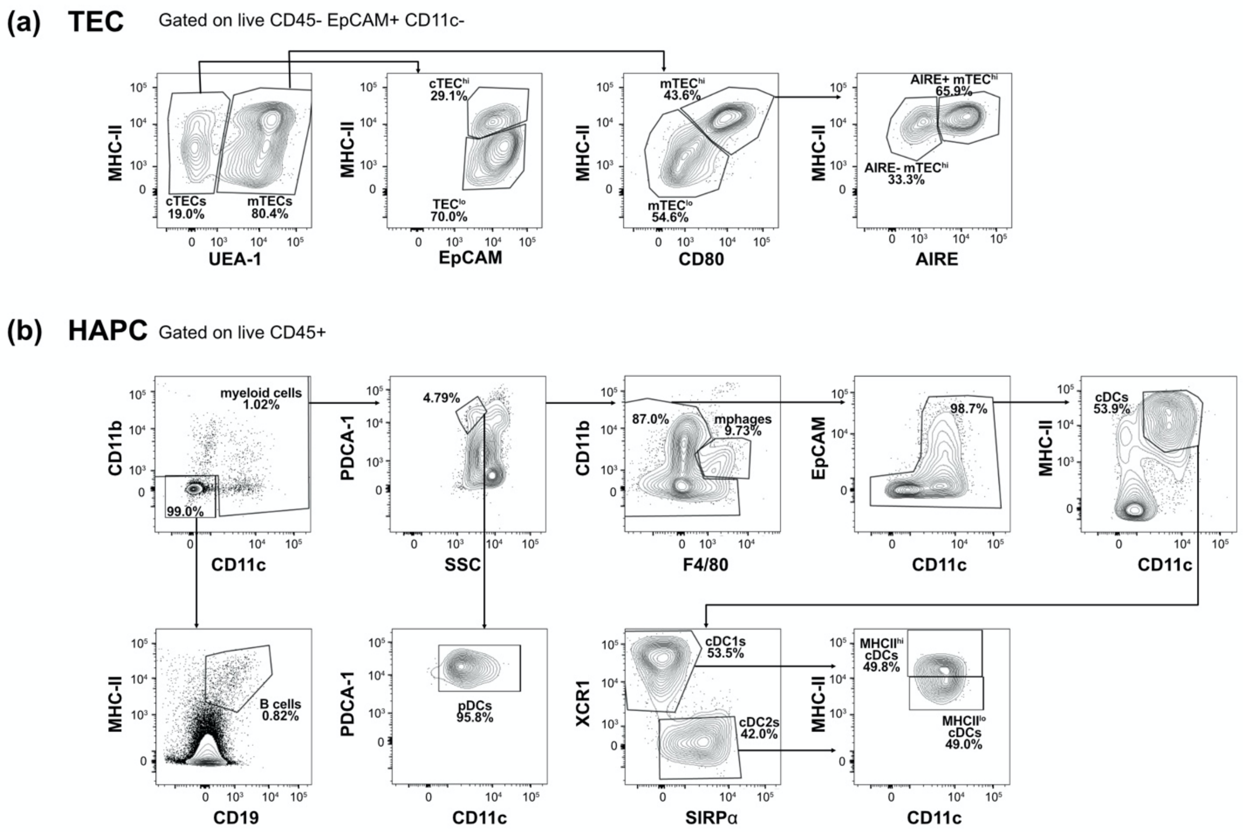


**Supplementary Figure 3, related to Figure 6. Flow cytometric analyses of TECs and HAPCs. (a)** Representative flow cytometry plots to quantify the frequency of cTEC and mTEC subsets within enzymatically digested 1MO and 12MO thymuses: TEC$^{lo}$ (CD45$^{-}$EpCAM$^{+}$CD11c$^{-}$UEA-1$^{-}$MHC-II$^{lo}$), cTEC$^{hi}$ (CD45$^{-}$EpCAM$^{+}$CD11c$^{-}$UEA-1$^{-}$MHC-II$^{hi}$), mTEC$^{lo}$ (CD45$^{-}$EpCAM$^{+}$CD11c$^{-}$UEA-1$^{+}$MHC-II$^{lo}$), AIRE$^{-}$ mTEC$^{hi}$ (CD45$^{-}$EpCAM$^{+}$CD11c$^{-}$UEA-1$^{+}$MHC-II$^{hi}$AIRE$^{-}$), AIRE$^{+}$ mTEC$^{hi}$ (CD45$^{-}$EpCAM$^{+}$CD11c$^{-}$UEA-1$^{+}$MHC-II$^{hi}$AIRE$^{+}$). **(b)** Representative flow cytometry plots to quantify the frequency of HAPC subsets within enzymatically digested 1MO and 12MO thymuses: B cell (CD45$^{+}$CD11b$^{-}$CD11c$^{-}$CD19$^{+}$MHC-II$^{+}$), pDC (CD45$^{+}$ PDCA-1$^{+}$CD11c$^{int}$SSC$^{lo}$), macrophages (CD45$^{+}$CD11b$^{+}$F4/80$^{+}$), cDC1 (CD45$^{+}$F4/80$^{-}$CD11c$^{+}$MHC-II$^{+}$XCR1$^{+}$), cDC2 (CD45$^{+}$ F4/80$^{-}$CD11c$^{+}$MHC-II$^{+}$SIRP$\alpha^{+}$). cDC1 and cDC2 subsets were further subdivided into MHC-II$^{lo}$ and MHC-II$^{hi}$ populations.

SUPPLEMENTARY MOVIE LEGENDS

**Supplementary Movie 1, related to Figure 1. Migration of 1MO and 12MO CD4SP thymocytes on a 1MO thymus slice.** 1MO (red) and 12MO (blue) CD4SPs migrate within an EGFP-expressing 1MO thymus slice (left panel). Thymocyte track times are color-encoded as indicated (right panel). Images were acquired for 15 min with 15 sec time intervals, through a depth of 40 μm, and a maximum intensity projection is displayed. Scale bar is 100 μm.

**Supplementary Movie 2, related to Figure 1. Migration of 1MO and 12MO CD4SP thymocytes on a 12MO thymus slice.** 1MO (red) and 12MO (blue) CD4SPs migrate within an EGFP-expressing 12MO thymus slice (left panel). Thymocyte track times are color-encoded as indicated (right panel). Images were acquired for 15 min with 15 sec time intervals, through a depth of 40 μm, and a maximum intensity projection is displayed. Scale bar is 100 μm.
